# Dual-domain Flower signaling coordinates extracellular vesicles-mediated fitness selection and cell-intrinsic survival in astrocytes

**DOI:** 10.1101/2025.08.08.669265

**Authors:** Szu-Min Tu, Ching-Hsin Lin, Yang Liu, Claudia Schirra, Chin-An Yang, Johannes Hohneck, Martin Jung, Annika Clemenz, Sandra Rother, Ahmad Aljohmani, Daniela Yildiz, Yvonne Schwarz, Elmar Krause, Walter J. Schulz-Schaeffer, Veit Flockerzi, Chi-Kuang Yao, Hsin-Fang Chang

## Abstract

Cellular fitness surveillance preserves tissue integrity, yet its regulation within the mammalian brain remains poorly understood. We identify a bifurcated mechanism in the transmembrane protein Flower—encoding both survival-promoting (“win”) and apoptosis-inducing (“lose”) isoforms. We demonstrate that astrocytes secrete specialized extracellular vesicles (EVs), termed “fitness vesicles,” carrying Flower to facilitate competitive selection across distal cell populations. The N-terminal Flower domain acts extrinsically via these EVs to drive the elimination of less-fit neighbors. Conversely, the “win“-specific C-terminal domain functions as a cell-intrinsic module; it translocates to the nucleus under stress to repress Caspase-3 and provide resilience. In Alzheimer’s disease (AD) models and human AD brains, Flower-positive astrocytes accumulate around amyloid-β (Aβ) plaques. Under Aβ stress, the “win” isoform reprograms astrocytes toward a neuroprotective state that enhances plaque clearance while ensuring cell-intrinsic survival. Our findings reveal how Flower couples long-range, EV-mediated cellular selection with cell-autonomous protection to coordinate astrocyte quality control and tissue resilience in neurodegeneration.

## INTRODUCTION

Intercellular communication is fundamental to brain homeostasis and occurs through both direct cell–cell contact and secreted extracellular vesicles (EVs)(Schnatz *et al*, 2021). Astrocyte-derived EVs modulate synaptic activity, inflammatory responses, and amyloid-β (Aβ) metabolism, underscoring their role in neural tissue regulation(Chaudhuri *et al*, 2018; Li *et al*, 2020b; Patel & Weaver, 2021; You *et al*, 2020). However, whether EV-mediated communication contributes to cellular fitness surveillance in the mammalian brain remains unknown. In particular, it is unclear whether molecular determinants of cellular fitness, conventionally viewed as mediators of contact-dependent competition, can be transmitted via EVs to regulate competitive cellular outcomes across extended distances beyond direct membrane contact.

The transmembrane protein Flower was first identified in *Drosophila melanogaster* as a regulator of synaptic vesicle endocytosis (Yao *et al*, 2009). Subsequent studies have confirmed Flower’s role in endocytosis in both invertebrates and vertebrates(Chang *et al*, 2018; Ravichandran *et al*, 2024; Xue *et al*, 2012; Yao *et al*., 2009; Yao *et al*, 2017). *Flower* is evolutionarily conserved from flies to mammals and is ubiquitously expressed across tissues, highlighting its ancient and essential biological functions. The gene encodes multiple isoforms, some of which are involved in calcium channel activity and vesicle trafficking(Brose & Neher, 2009; Kuo & Trussell, 2009; Li *et al*, 2020a; Seidenthal *et al*, 2025; Yao *et al*., 2009; Yao *et al*., 2017). More recently, specific Flower isoforms have been recognized as key regulators of a fundamental process known as cellular fitness competition(Merino *et al*, 2013; Rhiner *et al*, 2010), a natural selection mechanism that eliminates unfit cells by inducing apoptosis while allowing healthy, fit cells to continue growing(Madan *et al*, 2018; Merino *et al*, 2016; Merino *et al*, 2015). This selection process results in the survival of fitter ‘winner’ cells, while unfit ‘losers’ are eliminated from the population.

In heterogeneous cell populations, Flower isoforms function as molecular “fitness fingerprints” displayed on the cell surface(Merino *et al*., 2013; Rhiner *et al*., 2010). Fit cells predominantly express “win” isoforms, while less-fit cells express “lose” isoforms. This differential expression drives selective apoptosis of unfit cells and survival of fitter neighbors, establishing a winner–loser signaling axis. Such fitness-based cell selection is essential for tissue integrity during embryonic development, aging, infection, tumorigenesis, and neurodegeneration(Coelho & Moreno, 2019; Costa-Rodrigues *et al*, 2021; Gogna *et al*, 2015; Madan *et al*, 2020; Madan *et al*, 2019; Marques-Reis & Moreno, 2021; Parker *et al*, 2020; Yekelchyk *et al*, 2021). Despite its broad relevance, the mechanistic basis of Flower-mediated signaling in mammals remains poorly understood.

Tissue homeostasis depends on maintaining a balanced fitness landscape. Neurodegenerative conditions disrupt this equilibrium, generating heterogeneous cell populations with divergent survival capacities. Astrocytes are central regulators of brain homeostasis and play key roles in Aβ clearance and neuronal support in Alzheimer’s disease (AD)(Dai *et al*, 2023; Gomez-Arboledas *et al*, 2018; Jiwaji *et al*, 2022; Kim *et al*, 2022; Lines *et al*, 2022). However, astrocytic responses are highly heterogeneous(Moonen *et al*, 2023; Patani *et al*, 2023), suggesting the presence of intrinsic selection mechanisms that influence cell survival and function. Whether Flower-mediated fitness signaling contributes to this heterogeneity remains unknown.

Here, we present experimental evidence that points the existing model of Flower-mediated signaling in a new direction. Our topological analyses demonstrate that in intact astrocytes, both the N- and C-terminal signaling domains of Flower are localized in the cytoplasm, rendering them sequestered from direct intercellular contact. To resolve this paradox, we identify a novel extracellular vesicle (EV)–associated pathway that serves as the essential vehicle for signal presentation. We show that incorporation into specialized “fitness vesicles” triggers a topological transition, exposing these domains to the extracellular environment and enabling long-range intercellular selection across distal cell populations. Using complementary molecular and imaging approaches, we define the functional roles of this mechanism under Aβ stress, validating our findings in APP-transgenic mice and human Alzheimer’s disease tissue. Together, our results uncover a previously unrecognized dual-domain mechanism that bifurcates Flower signaling into EV-mediated distal selection and cell-intrinsic protection, providing a new model for understanding tissue resilience in neurodegeneration.

## RESULTS

### Flower isoforms mediate differential cell survival in primary astrocytes

Flower is a conserved cell fitness marker that labels cells as “winners” or “losers” during competitive interactions. In mice, the Flower locus encodes four isoforms (Fig. 1A), of which mFwe2 and mFwe4 are associated with a “win” phenotype, whereas mFwe1 and mFwe3 confer a “lose” identity(Petrova *et al*, 2012). To investigate how Flower isoforms regulate astrocyte fitness, we established a quantitative cell competition assay using primary astrocytes derived from Flower knockout (KO) mice.

**Figure 1.**
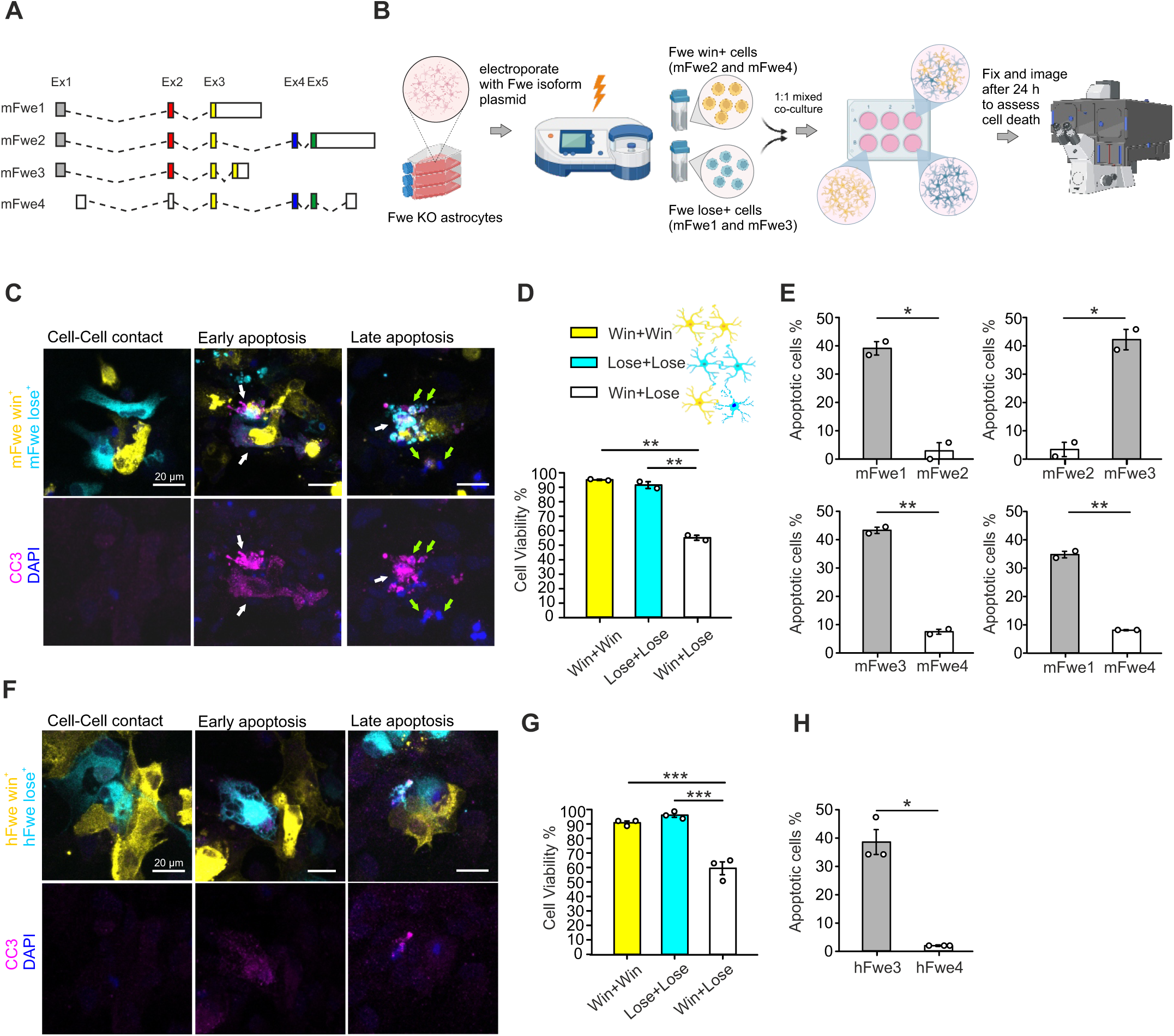
Flower-mediated regulation of cell fitness in primary astrocytes. **(A)** Schematic representation of mouse Flower isoforms (mFwe1–4), illustrating comparable exon and intron structures across splice variants. **(B)** Schematic of the experimental procedure. Primary astrocytes were isolated from Flower KO pups and cultured for 9–12 days before electroporation. Individual Flower isoforms were introduced into astrocytes via electroporation. Transfected cells were then mixed at a 1:1 ratio to establish homotypic and heterotypic co-cultures. After 24 h of co-culture in a 6-well plate, cells were fixed and stained for cell death markers. Confocal imaging was used to assess cell death resulting from Flower-mediated cell selection. **(C)** Confocal images of primary astrocytes expressing the “win” Flower isoform (yellow) co-cultured with astrocytes expressing the “lose” isoform (cyan). Cell-cell contacts between isoform-expressing cells were analysed after 24 h. Cultures were fixed and stained with cleaved caspase-3 (CC3; magenta) to mark apoptotic cells, while DAPI staining (blue), applied before fixation to identify non-intact cells or fragmented nuclei indicative of late apoptosis. Scale bar, 20 µm. White arrows indicate CC3+ apoptotic cells, and green arrows point to fragmented DAPI+ nuclei. **(D)** Quantification of cell death in homotypic pairwise win-win, lose-lose, and heterotypic win-lose co-cultures. Cell death was assessed based on CC3 and DAPI staining in each co-culture (n= 500-1200 cell contacts; N=3 independent mouse preparations). Each data point represents the mean of the indicated combinations: Win+Win (mFwe2+mFwe2, mFwe2+mFwe4, mFwe4+mFwe4), Lose+Lose (mFwe1+mFwe1, mFwe1+mFwe3), and Win+Lose (mFwe2+mFwe1, mFwe2+mFwe3, mFwe4+mFwe1, mFwe4+mFwe3). **(E)** Quantification of cell death in individual isoform-expressing cells in heterotypic win-lose pairwise co-cultures (n= 260-360 cell contacts; N=2 independent mouse preparations). **(F)** Confocal images of mouse primary Flower KO astrocytes expressing the human Flower “win” isoform 4 (yellow) co-cultured with astrocytes expressing the human “lose” isoform 3 (cyan). Cell-cell contacts between hFwe 3- and hFwe 4-expressing cells were analysed after 24 h. Cultures were fixed and stained with CC3 (magenta) to identify apoptotic cells. DAPI staining (blue), applied before fixation to define non-intact cells or fragmented nuclei indicative of late apoptosis. Scale bar, 20 µm. **(G)** Quantification of cell survival in co-cultures. Dead cells were counted based on CC3 and DAPI staining (N = 3 independent experiments; n= 430-760 cell contacts were analysed). Each data point represents the indicated combination: Win+Win (hFwe4 + hFwe4), Lose+Lose (hFwe3 + hFwe3), and Win+Lose (mean of reciprocal fluorophore labeling, hFwe4-mCherry + hFwe3-mTFP and hFwe4-mTFP + hFwe3-mCherry). Statistical significance was determined by one-way ANOVA followed by Holm-Sidak post hoc test for multiple comparisons (***p < 0.001). **(H)** Quantification of cell death in win-lose co-culture from **G**. Data are presented as mean ± SEM. Statistical significance was determined using Student’s t-test: *p < 0.05, **p < 0.01, ***p < 0.001.

Specific Flower isoforms were reintroduced into KO astrocytes by electroporation to generate defined winner and loser populations. Astrocytes expressing individual isoforms were then co-cultured in pairwise combinations, including homotypic (mFwe1 + mFwe1, mFwe2 + mFwe2, mFwe3 + mFwe3) and heterotypic (mFwe1 + mFwe2, mFwe1 + mFwe3, mFwe1 + mFwe4, mFwe2 + mFwe1, mFwe2 + mFwe3, mFwe2 + mFwe4, mFwe3 + mFwe4) conditions, enabling classification into win–win, lose–lose, and win–lose interactions (Fig. 1B).

After 24 h of co-culture, dynamic interactions were observed, particularly in heterotypic pairings, characterized by cell–cell contact and selective cell elimination. Cell death was assessed using two complementary approaches: pre-fixation application of the membrane-impermeable dye DAPI to label cells with compromised membrane integrity, and post-fixation immunostaining for cleaved caspase-3 (CC3) to detect apoptosis (Fig. 1C). Cell–cell contact pairs were analyzed to determine viability, with dead cells classified as either early apoptotic (CC3⁺) or late apoptotic/necrotic (DAPI⁺ with nuclear fragmentation).

In win–win and lose–lose homotypic cultures, overall cell death remained low (6–8%). In contrast, heterotypic co-cultures of win and lose isoforms exhibited a marked increase in apoptosis, with approximately 45% of cells undergoing cell death (Fig. 1D). Notably, apoptosis occurred predominantly in cells expressing the lose isoforms. In the Fwe1 + Fwe2, Fwe2 + Fwe3, Fwe3 + Fwe4, and Fwe1 + Fwe4 co-cultures, cells expressing either mFwe1 or mFwe3 were consistently eliminated, confirming their functional identity as “loser” isoforms. Overall, 35–45% of less-fit astrocytes were selectively removed through Flower-mediated cell competition (Fig. 1E).

To determine whether this fitness code is conserved across species, we next tested the human Flower isoforms hFWE3 (“lose”) and hFWE4 (“win”), whose phenotypes have been previously characterized in tumor models(Madan *et al*., 2019). As in the mouse system, homotypic win–win (hFWE4 + hFWE4) and lose–lose (hFWE3 + hFWE3) cultures showed minimal cell death. In contrast, co-culture of hFWE3-and hFWE4-expressing astrocytes resulted in ∼40% total cell death within the mixed population (Fig. 1F,G), with selective elimination of ∼40% of hFWE3-expressing cells (Fig. 1H).

Together, these results establish a robust primary astrocyte co-culture system that faithfully recapitulates Flower-dependent elimination of less-fit cells by fitter neighbors. This assay provides a quantitative platform for dissecting the molecular mechanisms and signaling domains underlying Flower-mediated fitness surveillance in glial cells.

### N- and C-termini of Flower isoforms localize to the cytoplasmic side in astrocytes

Cellular fitness is generally considered to be regulated via cell–cell contact(Rhiner *et al*., 2010), implying that the functional domains of Flower isoforms should be exposed on the cell surface. However, topological predictions for full-length mouse Flower isoforms are inconsistent with experimental observations(Chang *et al*., 2018; Petrova *et al*., 2012). To address this discrepancy, we first examined the expression of all Flower isoforms in primary mouse astrocytes by quantitative RT-PCR. All four mouse Flower isoforms were detected, with mFwe1 and mFwe2 representing the predominant lose and win isoforms, respectively (Fig. 2A). Based on their relative abundance and established functional relevance, we subsequently focused our topological analyses on mFwe1 (lose) and mFwe2 (win).

**Figure 2.**
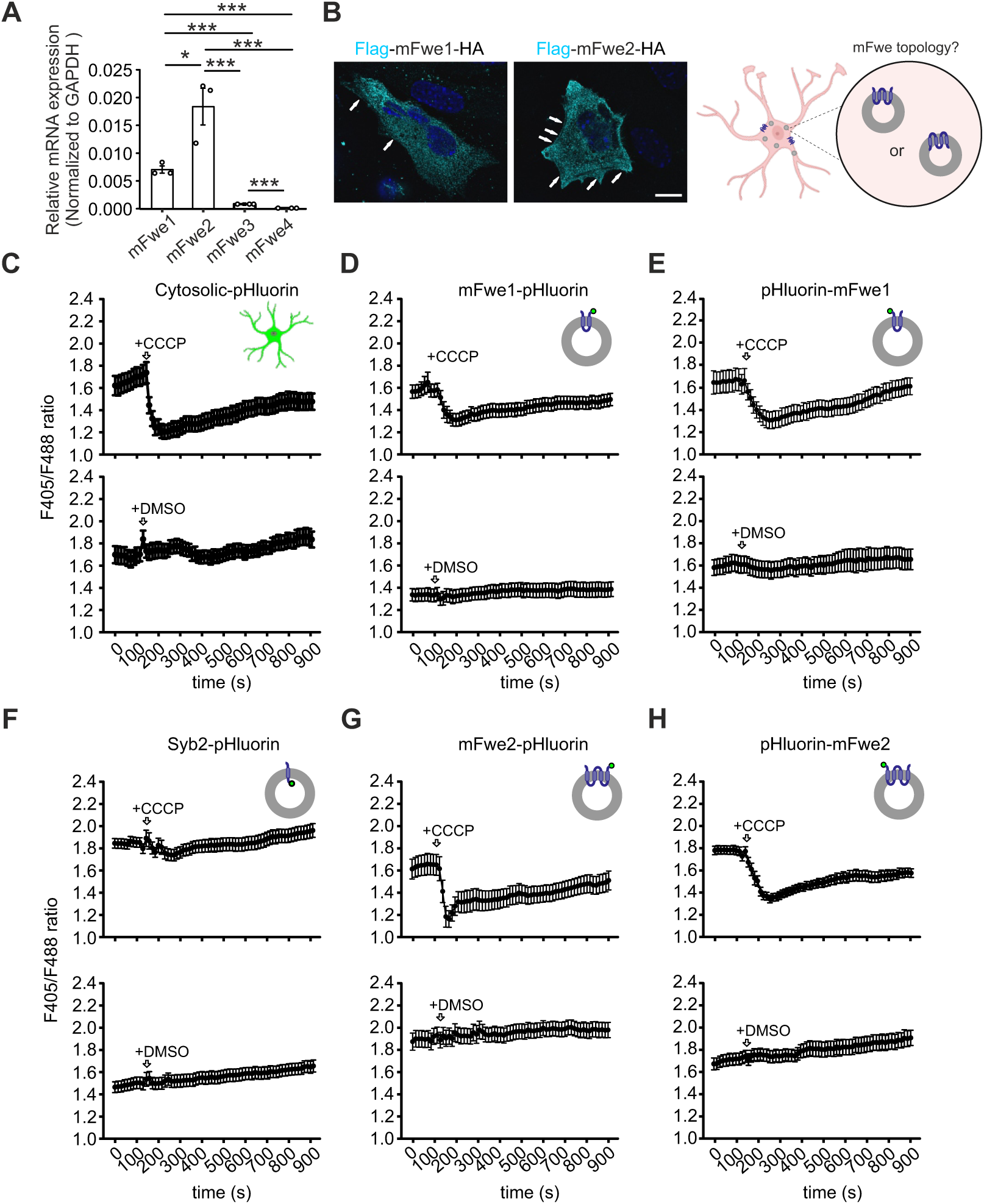
N- and C-termini of Flower proteins are localized on the intracellular side of mouse primary astrocytes. **(A)** Quantitative RT-PCR analysis of Flower mRNA expression of all Flower isoforms in WT astrocytes. (N = 3). Data are presented as mean ± SEM. Statistical significance was determined by one-way ANOVA followed by Holm-Sidak post hoc test for multiple comparisons: *p < 0.05 and ***p < 0.001. **(B)** Left: SIM images of WT astrocytes transfected with Flag–mFwe1–HA or Flag–mFwe2–HA. Cells were fixed and stained with anti-Flag antibody. White arrows indicate Flower localization at the plasma membrane. Scale bar: 10 µm. Right: Schematic illustrating the two potential membrane topologies of Flower on intracellular vesicles in astrocytes. **(C-H)** WT astrocytes were transfected with pH-sensitive pHluorin2 constructs to evaluate the orientation of the N- and C-termini of Flower proteins (mFwe1 and mFwe2) in relation to the vesicle lumen or cytoplasm. Cytosolic-pHluorin2 and Syb2-pHluorin2 were used as internal controls, representing known pHluorin2 localization in the cytoplasm and vesicle lumen, respectively. Ionophore CCCP was applied to transfected cells to induce pH changes, while DMSO served as the control vehicle. Live-cell confocal imaging was used to record ratiometric changes by sequential excitation at 405 nm and 488 nm. Construct details and sample sizes are as follows: cytosolic-pHluorin2 (DMSO, n = 19 cells; CCCP, n = 32 cells), mFwe1-pHluorin2 (DMSO, n = 34; CCCP, n = 57), pHluorin2-mFwe1 (DMSO, n = 29; CCCP, n = 31), Syb2-pHluorin2 (DMSO, n = 50; CCCP, n = 54), mFwe2-pHluorin2 (DMSO, n = 37; CCCP, n = 71), and pHluorin2-mFwe2 (DMSO, n = 42; CCCP, n = 60). N = 3 independent experiments.

Super-resolution SIM revealed that both Flag/HA tagged mFwe1- and mFwe2-expressing WT astrocytes displayed a punctate intracellular distribution, with partial localization at the plasma membrane, consistent with Flower being a vesicular membrane protein (Ravichandran *et al*., 2024; Yao *et al*., 2009) (Fig. 2B). To determine the topology of Flower isoforms, we fused the pH-sensitive GFP variant pHluorin2 to either the N- or C-terminus of mFwe1 and mFwe2. Primary astrocytes were transfected with these constructs and subjected to live-cell confocal imaging to monitor ratiometric fluorescence (405/488 nm) following cytosolic acidification induced by the protonophore carbonyl cyanide m-chlorophenyl hydrazone (CCCP). Astrocytes were maintained in DMEM during imaging, and DMSO-treated cells served as vehicle controls. Cytosolic pHluorin2 and vesicle-luminal Synaptobrevin2–pHluorin2 (Syb2-pHluorin2) were included as localization controls.

CCCP treatment induced a robust decrease in the 405/488 fluorescence ratio in cells expressing cytosolic pHluorin2, confirming effective cytoplasmic acidification (Fig. 2C). Both N- and C-terminally tagged mFwe1 constructs exhibited comparable decreases in fluorescence ratio (Fig. 2D,E), indicating cytoplasmic exposure of both termini. In contrast, vesicle-luminal Syb2-pHluorin2 remained unchanged despite robust cytosolic acidification (Fig. 2F), demonstrating that pHluorin2 confined to intracellular lumens is not responsive to CCCP-induced cytoplasmic pH shifts. Similarly, mFwe2 constructs tagged at either terminus displayed comparable decreases in fluorescence ratio upon CCCP treatment (Fig. 2G,H), indicating that both termini of mFwe2 are likewise cytoplasmic.

Because luminally oriented pHluorin2 does not respond to cytosolic acidification in this assay, Flower molecules with an inward-facing orientation would not contribute to the measured 405/488 ratio. The ratiometric signal therefore reflects the predominant cytoplasmic exposure of the tagged termini. Although the presence of a minor pool of Flower proteins with alternative orientation or subcellular localization cannot be excluded, the data indicate that the majority of both N- and C-termini of mFwe1 and mFwe2 are cytoplasmically exposed in astrocytes.

Together, these results establish that both the majority of lose isoform mFwe1 and the win isoform mFwe2 possess cytoplasmic N- and C-terminal domains in astrocytes. This topology challenges the prevailing model of Flower-mediated fitness regulation as a strictly contact-dependent process(Rhiner *et al*., 2010) and suggests that Flower may operate through an alternative, non–cell-autonomous signaling mechanism in astrocytes.

### Flower is released on extracellular vesicles to mediate fitness competition

In *Drosophila*, Flower-mediated fitness recognition relies on extracellular exposure of the C-terminal domain, enabling contact-dependent signaling between neighboring cells(Rhiner *et al*., 2010). In contrast, our topological analyses demonstrate that both N- and C-termini of Flower isoforms are oriented toward the cytoplasm in astrocytes, suggesting an alternative, non–cell-autonomous mechanism of signaling. To distinguish total Flower protein from isoform-specific C-terminal exposure, we employed two complementary antibodies: a polyclonal Pan-Flower antibody, raised against conserved regions and recognizing both mouse and human Flower proteins without distinguishing isoforms, and a custom C-terminal-specific antibody (#C1557) that selectively detects mouse win isoforms (mFwe2 and mFwe4) but recognizes both hFWE3 and hFWE4 in human tissue.

Consistent with the possibility of extracellular Flower, immunoreactivity detected with the #C1557 antibody was frequently observed plaque-adjacent regions in brain tissue from both APP-transgenic mice and Alzheimer’s disease patients (Fig. EV1A). While this extracellular signal did not overlap with astrocytic markers, its proximity to plaques suggests a secreted pool of Flower, potentially associated with EVs or amyloid-bound assemblies. To investigate the mechanism of release, we analyzed Flower processing in Flower KO astrocytes overexpressing Flag-Flower-HA constructs. Western blot analysis revealed the presence of both full-length Flower and lower molecular weight fragments detected by anti-Flag (N-terminal) and anti-HA (C-terminal) antibodies, consistent with partial proteolytic processing. Quantification was performed based on band intensity integration of full-length and cleaved fragments. Bands corresponding to lower molecular weight species within the same lane were considered processed forms. The win isoform mFwe2 exhibited substantial processing, with approximately 40% full-length protein and ∼60% cleaved fragments, whereas other isoforms remained predominantly full-length (70–80%) (Fig. EV1B–D).

We next asked whether Flower fragments are secreted via EVs. EVs were isolated from astrocyte culture supernatants by differential ultracentrifugation, with a final pelleting step at 100,000 × g. Western blot analysis of purified EV fractions from WT, mFwe1-, and mFwe2-overexpressing astrocytes showed CD63 enrichment, confirming existence of EV population (Fig. 3A). Immunostaining with the Pan-Flower antibody revealed Flower-positive puncta in supernatants from both WT and mFwe2-pHluorin2-expressing astrocytes (Fig. EV2A). SIM imaging showed that approximately 20% of these puncta co-localized with the membrane dye DiD, whereas ∼80% did not. This suggests that a majority of secreted Flower exists either in small, low-lipid vesicles below the DiD labeling threshold or as non-membranous protein-RNA/lipid complexes (Fig. EV2B).

**Figure 3.**
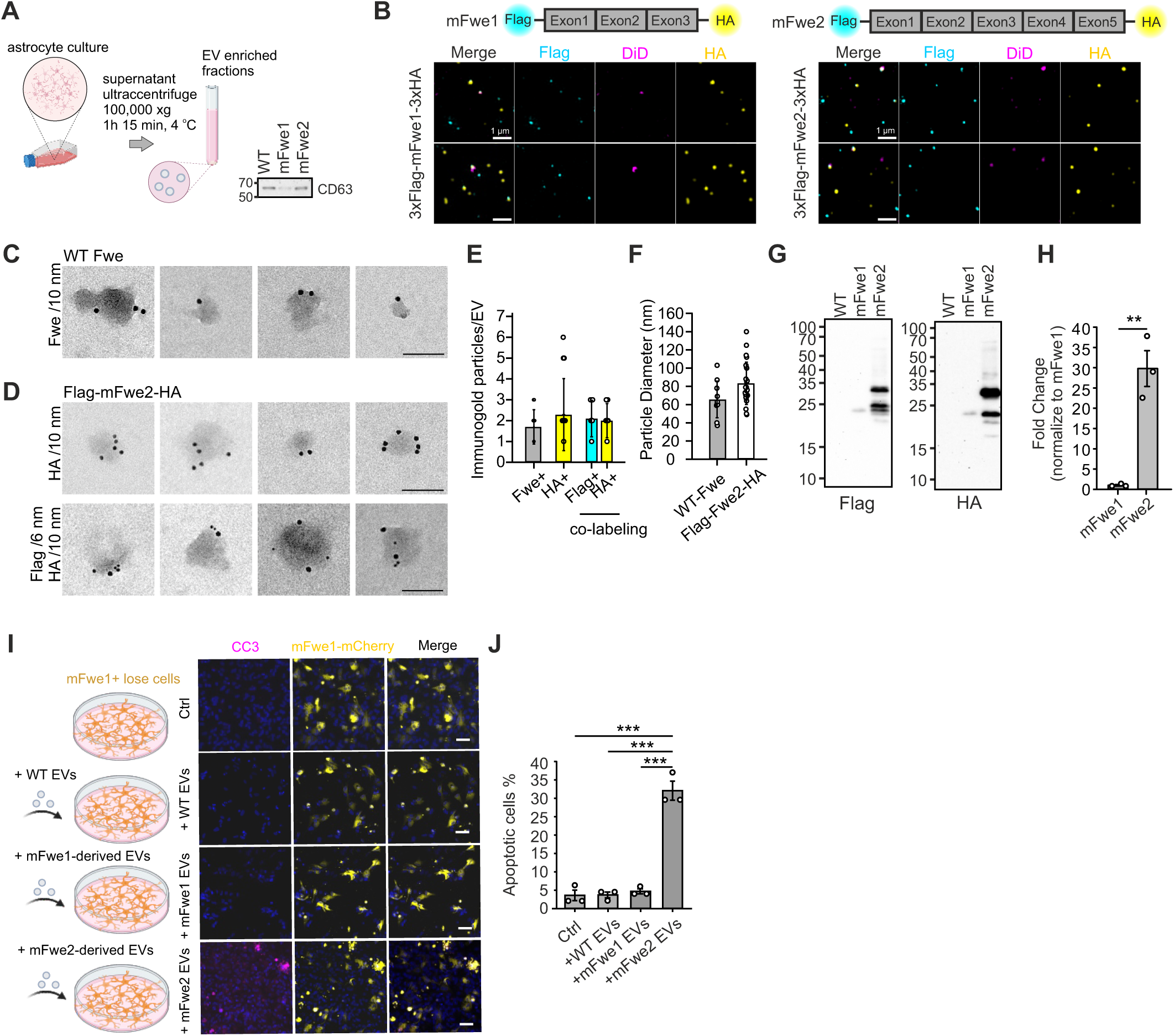
Extracellular vesicle–associated Flower mediates cellular fitness signaling. **(A)** Left: Experimental workflow for extracellular vesicle (EV) isolation. Astrocytes were cultured for two weeks following cortical isolation, and culture supernatants were collected for EV pelleting via ultracentrifugation. Right: Western blot analysis of EV lysates from WT astrocyte cultures transfected with mFwe1 or mFwe2. Ten micrograms of EV protein were loaded per lane. Anti-CD63 antibody was used to identify the EV population. **(B)** SIM images of EVs derived from astrocytes expressing Flag-mFwe1-HA or Flag-mFwe2-HA (right). Pelleted EVs were fixed and stained with anti-Flag (cyan), anti-HA (yellow), and DiD (magenta) to visualize N- and C-terminal fragments and their association with EV membranes. Schematic representations of Flower constructs (“win” Fwe2 and “lose” Fwe1) used in this study are shown (top). Scale bar, 1 μm. **(C)** Representative TEM images of EVs isolated from WT astrocyte culture supernatants. EVs were deposited onto pioloform-coated copper grids and immunogold-labeled with an anti-Pan-Flower primary antibody and a goat anti-rabbit secondary antibody conjugated to 10nm gold particles. Scale bar, 100 nm. (WT n_EVs_=10, N_Pups_=6). **(D)** TEM images of EVs isolated from WT astrocytes expressing Flag-mFwe2-HA. EVs were immunogold-labeled using anti-HA antibody and detected with a goat anti-rat secondary antibody conjugated to 10 nm gold particles (top). Co-labeling with anti-Flag and anti-HA antibodies was detected using goat anti-mouse (6 nm) and goat anti-rat (10 nm) gold-conjugated secondary antibodies, respectively (bottom). Scale bar, 100 nm. (anti-HA, n_EVs_=14; anti-Flag/anti-HA, n_Evs=_10; N_Pups_=6). **(E)** Quantification of gold particle numbers associated with immunolabeled EVs shown in **C** and **D**. Data are presented as mean ± SD. (WT n_EVs_=10; anti-HA, n_EVs_=14; anti-Flag/anti-HA, n_Evs=_10; N_Pups_=6). **(F)** Size distribution analysis of immunogold-labeled EVs shown in **C** and **D**. Data are presented as mean ± SD. (WT n_EVs_=10, N_Pups_=6). **(G)** Western blot analysis of secreted mFwe1 and mFwe2 in EV fractions isolated from WT astrocytes transfected with Flower constructs. Ten micrograms of EV protein were loaded per lane. Membranes were sequentially probed with anti-Flag and anti-HA. **(H)** Quantification of relative levels of secreted Fwe2 compared to Fwe1 from (**G)** (N = 3 independent experiments). Data are presented as mean ± SD. Statistical significance was determined using Student’s t-test: **p < 0.01. **(I)** Experimental workflow assessing the effect of Flower-containing EVs on cellular fitness (left). EVs isolated from WT, mFwe1-, or mFwe2-expressing astrocytes were applied to unfit cell populations. Representative confocal images of mFwe1-expressing unfit cells treated with EVs derived from WT, mFwe1-, or mFwe2-overexpressing astrocytes (right). Approximately 10 μg of EVs were applied to 100,000 recipient cells seeded on 12.5-mm coverslips in 24-well plates. After 24 h, cells were fixed and stained with anti-CC3 to label apoptotic cells. Scale bar, 50 μm. **(J)** Quantification of apoptotic cells from (H) (control n = 616; WT EVs n = 665; mFwe1 EVs n = 687; mFwe2 EVs n = 917; N = 3). Data are presented as mean ± SEM. Statistical significance: one-way ANOVA with Tukey’s post hoc test, *p < 0.001.

To assess isoform-specific secretion, we analyzed supernatants from astrocytes expressing either the win isoform mFwe2 or the lose isoform mFwe1, each tagged with Flag and HA at the N- and C-termini, respectively (Fig. 3B). SIM imaging of supernatants from astrocytes expressing N- and C-terminally tagged mFwe1 or mFwe2 revealed double-positive (Flag⁺/HA⁺) as well as single-positive puncta. These observations indicate the presence of Flower isoforms containing both termini, as well as puncta associated with isolated N- or C-terminal domains (Fig. EV2C). Particle size analysis showed comparable EV radii across conditions (mean radius: 71.6 nm for GFP control, 74.9 nm for mFwe1, and 76.0 nm for mFwe2; Fig. EV2D). In this experiment, mFwe1-expressing astrocytes released 3.7 × 10⁷ particles per million cells, while mFwe2-expressing cells released 1.6 × 10⁷ particles per million cells, indicating detectable EV secretion from both isoforms.

Transmission electron microscopy (TEM) combined with immunogold labeling provided direct ultrastructural evidence that Flower is present on secreted EVs. In WT astrocyte-derived EVs, 10 nm gold particles conjugated to the Pan-Flower antibody were detected on the surface of morphologically intact vesicles (Fig. 3C), establishing extracellular localization of endogenous Flower. In EVs derived from Flag-mFwe2-HA–expressing astrocytes, both 6 nm (Flag) and 10 nm (HA) gold particles were detected on single vesicles (Fig. 3D), demonstrating that full-length mFwe2 is incorporated into EV membranes with extracellular exposure of both termini. Controls without primary antibody showed minimal background (Fig. EV2E). Across conditions, EVs carried an average of ∼2 Flower molecules per vesicle, with mean diameters of ∼65 nm for WT EVs and ∼83 nm for mFwe2-overexpressing EVs (Fig. 3E,F).

Western blot analysis of purified EV fractions confirmed the incorporation of mFwe1 and mFwe2 into EVs, as detected by anti-Flag, anti-HA, and Pan-Flower antibodies. The absence of GAPDH in the EV fractions confirmed the purity of the preparation and the absence of cellular debris (Fig. 3G; Fig. EV2F). Despite equal loading (10 µg EV protein), endogenous Flower was below detection level, whereas mFwe2 consistently produced strong signals. Blots were initially probed with anti-Flower, then stripped and sequentially reprobed with anti-HA and anti-Flag antibodies. For quantitative comparison of isoform-specific secretion into EVs, anti-Flag signals were used because they exhibited high specificity and a linear detection range under the experimental conditions. Signal intensities were normalized to the number of transfected cells. Astrocytes expressing mFwe2 secreted ∼25-fold more Flower into EVs than those expressing mFwe1, revealing pronounced isoform-specific differences in EV release (Fig. 3H). Notably, although mFwe1-overexpressing cells produced more particles than mFwe2-expressing cells, the total Flower protein carried by mFwe2-derived EVs was substantially higher. Differences in transfection efficiency were modest (51% for mFwe2 vs. 42% for mFwe1), and mean fluorescence intensity per cell was comparable, indicating similar expression levels on a per-cell basis (Fig. EV2G,H).

Finally, we assessed the functional capacity of these vesicles to execute fitness selection. Treatment of ‘unfit’ (mFwe1-expressing) astrocytes with EVs purified from mFwe2-expressing cells induced robust apoptosis (>30%), whereas WT or mFwe1-derived EVs failed to elicit a death response (Fig. 3I,J). These data establish that EV-associated mFwe2 is a functionally potent effector of fitness competition. Together, our results uncover a non–cell-autonomous mechanism of Flower signaling that is fundamentally distinct from the classical contact-dependent model, providing a framework for long-range fitness surveillance in the mammalian brain.

### Flower is secreted via a non-canonical, CD63-independent vesicle population

To characterize the biogenesis of Flower-positive EVs, we compared their molecular signature to canonical exosomes using a topologically-defined CD63-pHuji reporter (Verweij *et al*, 2018). Intracellular analysis in astrocytes revealed that both Flower isoforms (mFwe1 and mFwe2) showed minimal co-localization with CD63-positive multivesicular bodies (MVBs) (Fig. EV3A,B). This suggests that the Flower protein bypasses the classical endosomal sorting pathway.

Super-resolution SIM imaging and ultrastructural analysis of isolated EVs from mFwe2 overexpressed astrocytes confirmed that Flower and CD63 reside in largely distinct vesicle populations. While a minor fraction (up to 15%) of CD63⁺ exosomes contained sequestered Flower within their lumen, the vast majority (∼70%) of Flower-positive puncta were CD63-negative (Fig. EV3C,D). Immunogold TEM further demonstrated that on these CD63-negative vesicles, both the N- and C-termini of Flower are exposed on the external surface (Fig. EV3E,F). The specificity of this labeling was confirmed using secondary-antibody-only controls, which showed minimal gold particle background (Fig. EV3G).

This “extracellular-out” topology is incompatible with the inward budding mechanism of MVB-derived exosomes, which would instead sequester the termini inside the vesicle lumen. These data establish that Flower is predominantly secreted via a non-canonical pathway, likely involving direct budding or recruitment to the limiting membrane of a specialized vesicle class. Based on their distinct molecular identity, unique topology, and role in fitness signaling, we designate these CD63-independent, Flower-exposing vesicles as “fitness vesicles” (Fig. EV3H).

### Flower’s N-terminus induces apoptosis, whereas its C-terminus antagonizes death signaling

To identify functional regions of Flower involved in fitness signaling, we focused on three domains containing potential interaction motifs: the N-terminus (YxxL)(Liu *et al*, 2012), exon 3 (conferring a win phenotype in human cells)(Madan *et al*., 2019), and the C-terminus (YxxI)(Cheng *et al*, 2017) (Fig. 4A). Our findings demonstrate that while the N- and C-terminal domains are oriented toward the cytoplasm, they are exposed on the outer membrane of astrocyte-derived EVs. To enrich these signaling domains on the cell surface, we engineered them to face the outer plasma membrane. Given that Flower functions as a fitness code(Rhiner *et al*., 2010), we hypothesized that these specific domains mediate the intercellular signals regulating survival or apoptosis. To test this, we generated truncated Flower constructs fused to mScarlet I and a platelet-derived growth factor receptor (PDGFR) transmembrane domain (mFwe-mScarlet I-TM), allowing ectopic presentation of individual Flower domains at the plasma membrane (Fig. EV4A). Surface localization of these constructs in astrocytes was confirmed by RFP and HA staining under non-permeabilizing conditions (Fig. EV4B). The constructs were expressed in Flower KO astrocytes, which were then separately co-cultured with either mFwe2-expressing (fit) or mFwe1-expressing (unfit) cells to assess cell death (Fig. 4B).

**Figure 4.**
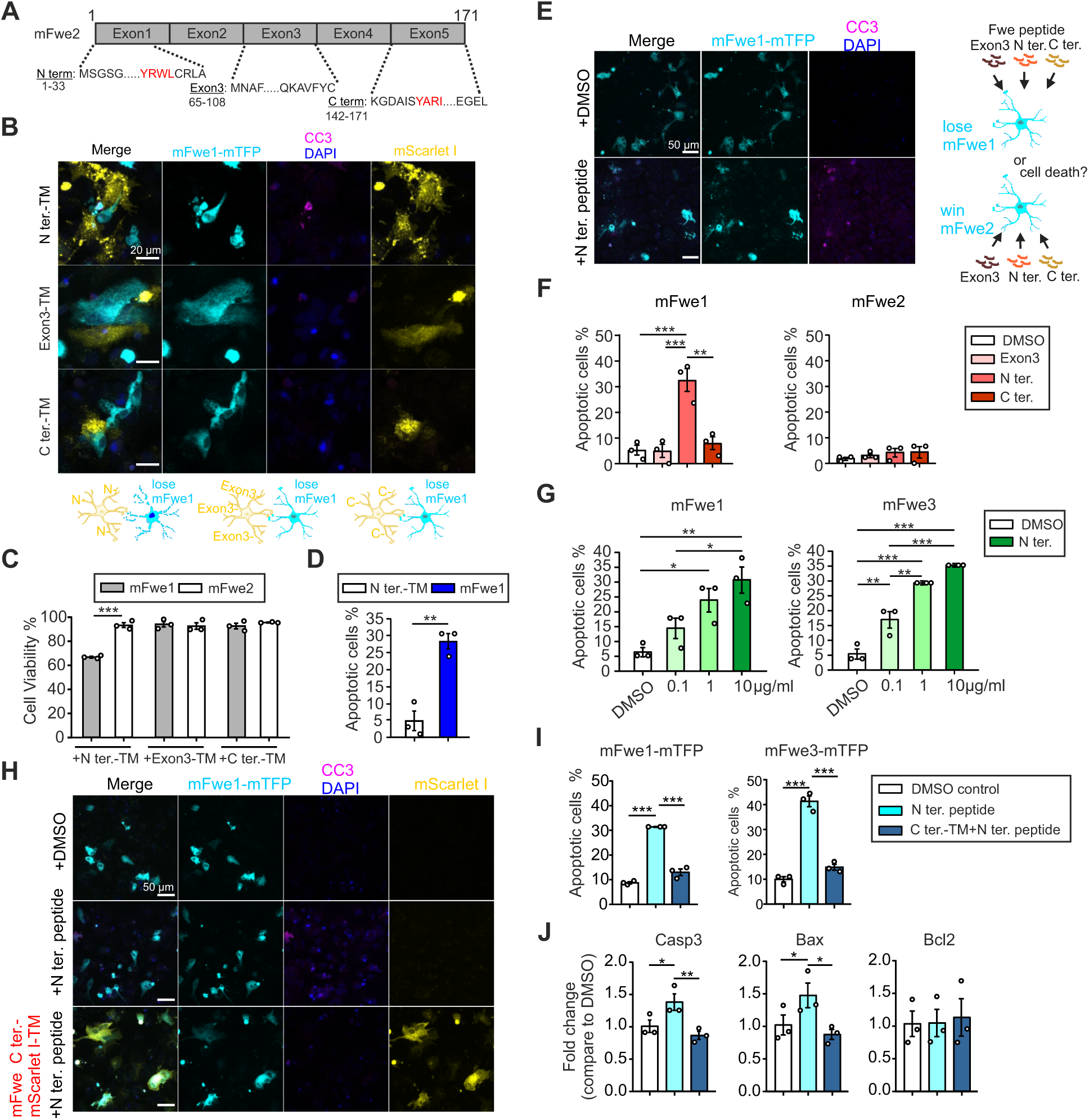
The Flower N-terminus acts as an extrinsic pro-apoptotic ligand antagonized by the C-terminal domain. **(A)** Schematic representation of putative signaling domains located at the N-terminus, C-terminus, and Exon 3 region of the full-length mFwe2 sequence. **(B-J)** Cell fitness assays were performed using Flower KO astrocytes expressing the indicated Flower isoforms to generate fit or unfit cell populations. Cells were treated with synthesized Flower N-terminal, C-terminal, or exon 3-derived peptides, or co-overexpressed with the corresponding truncated Flower constructs for cell-death rescue experiments. Cultures were fixed 24 h after treatment or co-culture, and cell death was assessed based on DAPI staining or CC3+ immunostaining. **(B)** Confocal images of cells expressing specific Flower signaling domains (yellow) localized at the plasma membrane (-TM) with extracellular orientation, co-cultured with unfit mFwe1-mTFP-expressing “lose” cell populations (cyan). Cultures were stained with DAPI before fixation and anti-CC3 antibody (magenta) after fixation to analyse positively stained dead cells. Scale bar, 20 µm. **(C)** Quantification of overall cell viability in co-cultures of Flower KO astrocytes expressing individual signaling domains (N terminus-, exon3- or C terminus-TM) with either fit (mFwe2) or unfit (mFwe1) cell populations (n = 255–342 cells per group; N = 3 independent experiments). Statistical significance was determined by two-way ANOVA followed by Holm-Sidak post hoc test for multiple comparisons: ***p < 0.001. **(D)** Detailed analysis of cell death in co-cultures of KO astrocytes expressing the N-terminal Flower construct (Flower N-term-TM) and mFwe1-expressing unfit cells, as shown in the first group of panel (**C**) (n = 307 cells; N = 3 independent experiments). Statistical significance was determined using Student’s t-test: **p < 0.01. **(E)** Confocal images of synthetic Flower N-terminus peptides applied to mFwe1-expressing unfit cells. Cultures were fixed and stained as described. Control cells were treated with DMSO. Scale bar, 50 µm. **(F)** Synthetic Flower signal peptides affect cell fitness. Peptides representing three Flower domains were incubated with mFwe1- or mFwe2-expressing cells for 24 h, and cell death was quantified. Data represent >300 cells per condition from N = 3 independent experiments. Statistical significance was determined by one-way ANOVA followed by Tukey’s post hoc test (**p < 0.01, ***p < 0.001). **(G)** Dose-dependent effect of the Flower N-terminal peptide on apoptosis in unfit cells. mFwe1- and mFwe3-expressing cells were treated with 0.1, 1, or 10 μg/ml of N-terminal peptide or DMSO control for 24 h. Cell death was quantified (n = 311–631 cells per condition; N = 3 independent experiments). Statistical significance was determined by one-way ANOVA with Tukey’s post hoc test (*p < 0.05, **p < 0.01, ***p < 0.001). **(H-I)** Role of the Flower C-terminus-TM peptide in rescuing cell death induced by the N-terminus peptide. mFwe1- and mFwe3-expressing unfit cells were treated with suspended Flower N-terminus peptide and transfected with or without the C terminus-TM construct. **(H)** Representative confocal images of mFwe1-expressing unfit astrocytes co-expressing the C-terminal-TM construct and treated with synthetic N-terminal peptide. Unfit cells without C-terminal-TM expression, treated with DMSO or N-terminal peptide, served as controls. Cells were fixed and stained with CC3 antibody to label apoptotic cells. Scale bar: 50 µm. **(I)** Quantification of dead cells following N-terminal peptide treatment, with or without co-expression of the C-terminal domain. mFwe1- and mFwe3-expressing cells were treated for 24 h, and cell death was quantified (n = 278–617 cells per condition; N = 3 independent experiments). Statistical significance was determined by two-way ANOVA followed by Holm-Sidak post hoc test for multiple comparisons: ***p < 0.001. **(J)** Quantitative RT-PCR analysis of Flower signal peptide-mediated cell death. KO astrocytes were transfected with mFwe1 to generate unfit cells and treated with the N-terminus peptide, with or without rescue by overexpression of Flower C- or N-terminus constructs. Apoptosis-related genes (Caspase-3, Bax, and Bcl2) were analysed after 24 h (N = 3). Data are presented as mean ± SEM. Statistical significance was determined by two-way ANOVA followed by Holm-Sidak post hoc test for multiple comparisons: *p < 0.05 and **p < 0.01.

Expression of the N-terminal-TM Flower domain selectively induced apoptosis in unfit cells when co-cultured with fit cells, whereas fit cells remained unaffected (Fig. 4C,D; Fig. EV4C). To determine whether this effect was mediated directly by the Flower peptide rather than by secondary cellular interactions, we treated cells with synthetic peptides corresponding to the truncated Flower domains (Fig. 4E). Consistent with the co-culture experiments, the synthetic N-terminal peptide induced apoptosis specifically in unfit cells (Fig. 4F). This effect was dose-dependent, with apoptosis reaching approximately 35% at higher peptide concentrations, closely matching the magnitude observed in the cell-based assays (Fig. 4G).

Given that win isoforms of Flower contain a unique C-terminal domain, we next tested whether this region confers resistance to N-terminal-mediated apoptosis. Unfit cells expressing mFwe1 or mFwe3 were co-transfected with a C-terminal-TM construct and exposed to the N-terminal peptide. Expression of the C-terminal domain significantly reduced apoptosis under these conditions (Fig. 4H,I). Importantly, overexpression of the C-terminal domain alone did not enhance basal cell survival (Fig. EV5A,B), indicating that this domain does not actively promote survival but instead antagonizes N-terminal–induced death signaling.

To further define the downstream pathways involved, we analyzed the expression of apoptosis-related genes in unfit cells treated with the N-terminal peptide, with or without C-terminal-TM domain expression. N-terminal peptide treatment induced the upregulation of *Caspase3* and *Bax* mRNA, consistent with activation of the intrinsic apoptotic pathway. Conversely, co-expression of the C-terminal domain suppressed this induction, while *Bcl2* levels remained stable (Fig. 4J). These results demonstrate that Flower-mediated death primarily engages a transcriptional *Bax-Caspase3* axis.

Together, these results demonstrate that Flower’s N-terminus triggers apoptosis in unfit astrocytes, whereas the C-terminus counteracts this death signal.

### Direct interaction between Flower N- and C-terminal domains modulates pro-apoptotic fitness signaling

Our functional assays (Fig. 4) identified a fundamental dichotomy in Flower signaling: the N-terminal domain triggers apoptosis in “unfit” astrocytes, while the C-terminal domain antagonizes this death signal. Since the N- and C-termini of Flower are cytoplasmically oriented in astrocytes, this suggests that their antagonistic interaction is specifically coordinated upon secretion.

To determine the biochemical basis for this antagonism, we performed Surface Plasmon Resonance (SPR) assays using synthetic peptides. A C-terminal peptide was immobilized on the sensor chip, and N- or C-terminal peptides were injected as analytes. SPR revealed a specific, concentration-dependent interaction between the N- and C-terminal peptides, whereas no self-association was observed for the C-terminal peptide alone (Fig. EV6A,B). Notably, the N-terminal peptide exhibited strong dose-dependent self-association (Fig. EV6C,D), suggesting that it may oligomerize to enhance its pro-apoptotic signaling potency.

These findings provide a biochemical framework for the modular signaling logic of Flower. We propose that the extracellular C-terminal domain attenuates death signaling through direct binding-dependent interference, effectively masking the N-terminal “death ligand“

Based on these results, we propose a model for Flower-mediated fitness regulation (Fig. 5). In the intracellular environment, Flower termini are cytoplasmically sequestered. Upon release via “fitness vesicles,” the preservation of membrane topology results in an “extracellular-out” orientation, exposing both domains on the EV surface. In this extracellular context, the N-terminal fragment transmits pro-apoptotic signals to promote the elimination of distal unfit cells. However, on “win” (mFwe2) vesicles or target cell surfaces, the C-terminal domain acts as a molecular shield, neutralizing the N-terminal signal through direct physical binding.

**Figure 5.**
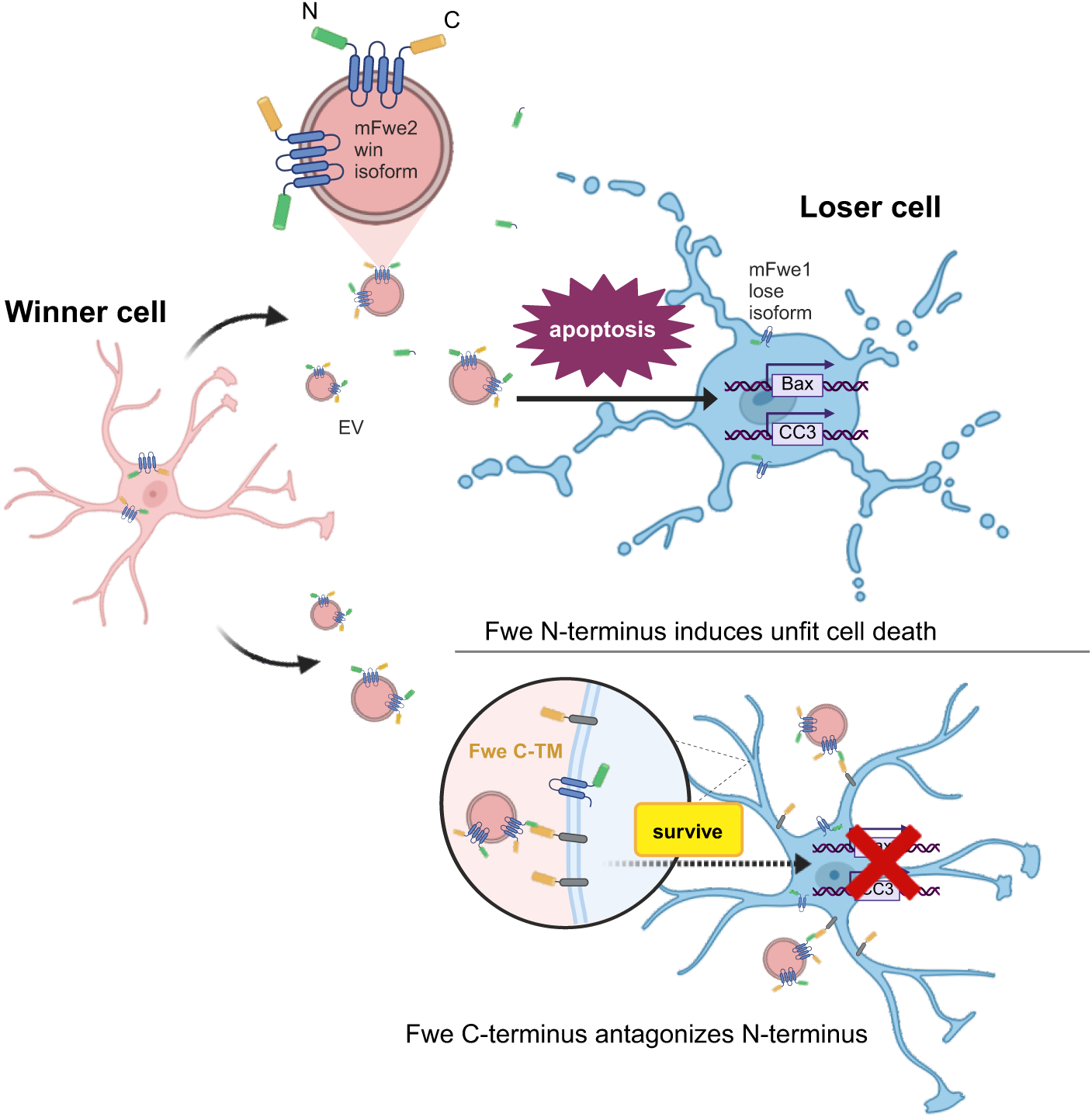
Model of EV-mediated astrocyte fitness surveillance and distal cell elimination. “Win” (mFwe2) astrocytes secrete specialized extracellular vesicles (EVs) displaying N-terminal “death” domains in an “extracellular-out” orientation. These EVs act as long-range signals that target “lose” (mFwe1) astrocytes, triggering a pro-apoptotic transcriptional program defined by *Bax* and *Caspase-3* induction. In contrast, the C-terminal domain (Fwe C-TM) functions as a competitive antagonist. By directly binding to the EV-derived N-terminal domain, the C-terminus physically neutralizes the pro-apoptotic signal. This binding-dependent interference protects “unfit” astrocytes from elimination.

Consistent with this framework, both “win” (mFwe2) and “lose” (mFwe1) isoforms contain the pro-apoptotic N-terminal region. However, only the mFwe2 isoform includes the C-terminal antagonistic domain, enabling the selective protection of “fit” cells from fitness-induced elimination through this localized, surface-masking mechanism.

### Localized enrichment of Flower isoforms in astrocytes surrounding amyloid plaques in mouse and human Alzheimer’s disease brains

To investigate whether Flower-mediated fitness signaling is engaged in human neurodegeneration, we first analyzed the expression of the Flower gene *CACFD1* in hippocampal specimens from patients with Alzheimer’s disease (AD) and age-matched healthy controls. Analysis of publicly available microarray data revealed that *CACFD1* expression positively correlates with AD disease severity, including incipient, moderate, and severe stages, as assessed by Braak stage and neurofibrillary tangle (NFT) score (Blalock *et al*, 2004) (Fig. 6A). While control individuals exhibited generally low *CACFD1* expression, transcript levels progressively increased with advancing disease stage. This gradual upregulation resulted in a significant elevation of *CACFD1* expression in severe AD patients compared with healthy donors (Fig. 6B), indicating an association between Flower expression and disease progression in the human brain.

**Figure 6.**
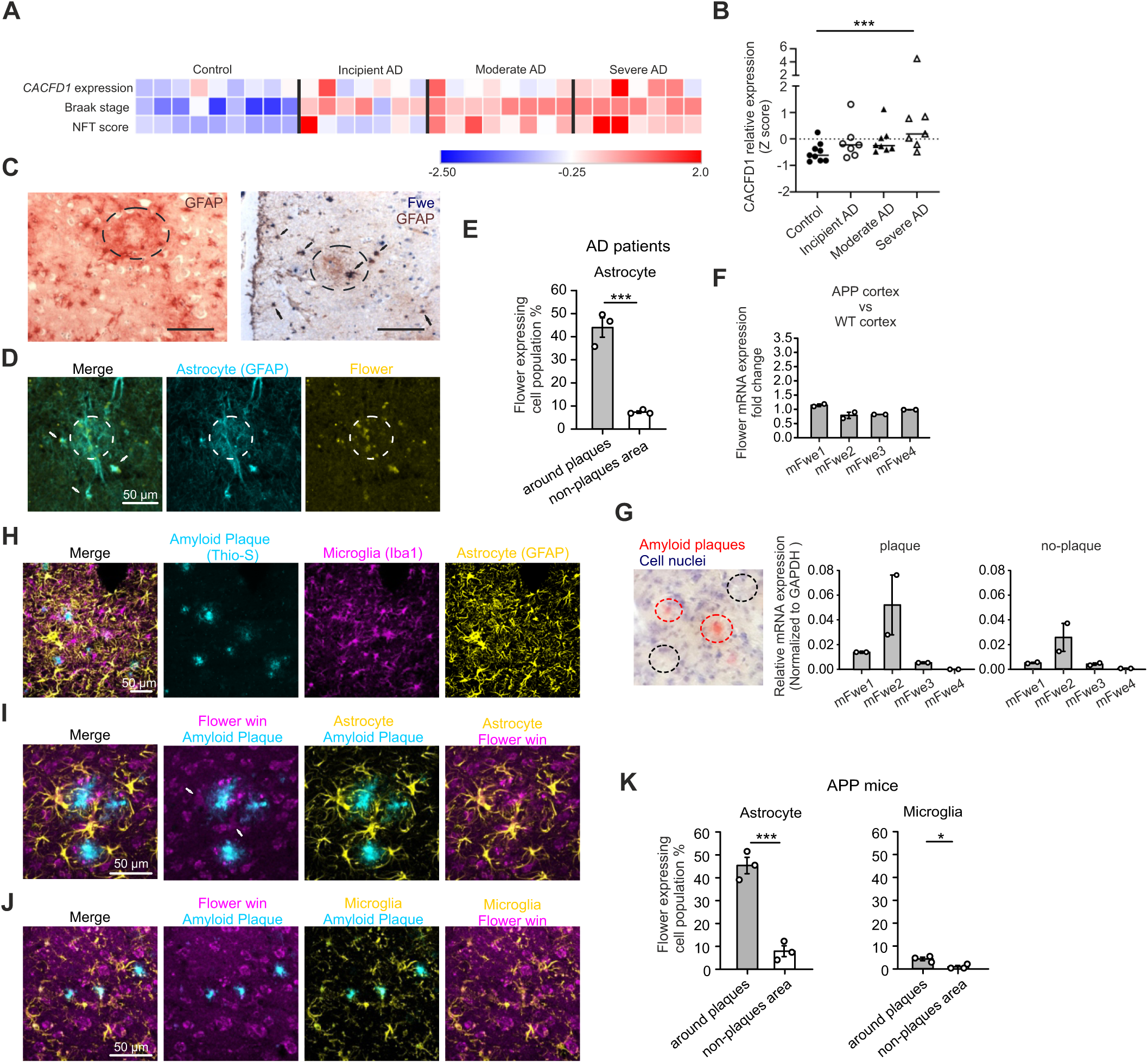
Flower is highly expressed in a subset of astrocytes near amyloid plaques in the neocortex. **(A)** Analysis of microarray data from hippocampal specimens of AD patients with different disease severities and from controls (GSE1297)(Blalock *et al*., 2004). Heatmap showing the correlation between CACFD1 expression and Braak stage or neurofibrillary tangle (NFT) score. Values are normalized to z-scores (sample value minus dataset mean divided by standard deviation). The heatmap was generated using Morpheus (https://software.broadinstitute.org/morpheus/); the color scale represents z-scores. **(B)** Relative CACFD1 expression across diagnostic groups. Lines indicate medians. P < 0.05 by two-tailed Kruskal–Wallis test. Control, n = 9; incipient AD, n = 7; moderate AD, n = 8; severe AD, n = 7. **(C)** Histological staining of post-mortem brain cortex from AD patients. Plaques are marked with black circles. Left: astrocytes stained with anti-GFAP antibody. Right: co-staining for Flower (blue) and astrocytes (brown). Black arrows indicate double-positive cells. Scale bar: 50 µm. **(D)** Confocal images of post-mortem brain cortex from AD patients. Flower (yellow) and GFAP (cyan) were stained to label Flower-expressing astrocytes. White arrows point to Flower-expressing astrocytes near plaques. **(E)** Quantification of Flower-expressing astrocyte populations near plaques versus non-plaque areas from (**D**) (N = 3 donor; n = 45 images; n = 1096 cells). **(F)** RT–qPCR analysis of total Flower expression in cortices from APP transgenic mice compared with WT controls. Data are presented as fold change relative to WT cortex. Flower isoform transcripts were normalized to GAPDH. Flower isoforms were normalized to GAPDH (N = 2). **(G)** Left: Representative cortical section from an APP mouse stained with Congo Red to label amyloid plaques (red) and hematoxylin to label nuclei (blue). Plaque regions (red circles) and non-plaque regions (black circles) were microdissected for RT–qPCR analysis. Right: RT–qPCR analysis of Flower isoform expression in plaque- or non-plaque-associated regions of APP mouse cortex (N = 2). **(H-J)** Representative confocal images of the neocortex from APP mice. Amyloid plaques were labeled with Thioflavin-S (Thio-S) (cyan), and glial cells were stained with anti-GFAP (astrocytes) and anti-IBA1 (microglia). Scale bar: 50 µm. **(I-J)** Endogenous Flower was stained with anti-Flower antibody #C-1557 (magenta) along with glial markers (yellow). White arrows indicate Flower-expressing astrocytes near plaques. **(K)** Quantification of Flower-high-expressing astrocytes and microglia in plaque-associated regions compared to non-plaque areas, based on the total counted glial cells. Left: astrocytes; right: microglia, derived from images in (**I**) and (**J**) (N = 3 mice; n = 30 images; 1167 astrocytes and 967 microglia were analysed). Data are presented as mean ± SEM. Statistical significance was determined using Student’s t-test: *p < 0.05, ***p < 0.001.

To validate these transcriptomic findings at the cellular level, we examined post-mortem cortical tissue from AD patients by immunohistochemistry. Consistent with previous reports, GFAP⁺ astrocytes accumulated around amyloid plaques(Sanchez-Mico *et al*, 2021). Notably, a subset of these plaque-associated astrocytes—defined as cells located within approximately 50 µm of plaque borders—displayed robust Flower immunoreactivity (Fig. 6C,D). Quantitative analysis revealed that ∼45% of astrocytes in plaque-associated regions expressed high levels of Flower, compared with ∼7% in non-plaque areas (Fig. 6E). The pronounced enrichment of Flower-positive astrocytes near plaques indicates locally elevated Flower signaling within the plaque microenvironment in human AD brains.

To determine whether Flower-mediated fitness signaling is engaged in vivo during neurodegeneration, we analyzed the APP transgenic mouse model of Alzheimer’s disease, which develops amyloid plaque pathology. We hypothesized that plaque-associated cortical regions may represent microenvironments of heterogeneous cellular fitness regulated by Flower-dependent signaling. RT–qPCR analysis of total cortical tissue showed no significant difference in overall Flower isoform expression between APP and WT mice (Fig. 6F). To examine spatial regulation, plaque-adjacent and plaque-free cortical regions were isolated using laser capture microdissection following Congo Red staining. Quantitative RT–PCR analysis showed that mFwe2 is the predominant Flower isoform in both plaque-associated and non-plaque cortical regions (Fig. 6G). However, mFwe2 expression was higher in plaque-adjacent regions compared with plaque-free cortical tissue, suggesting spatially restricted regulation of Flower expression during amyloid pathology.

To selectively assess Flower win isoform expression in mouse tissue, we performed immunostaining using the win-isoform–specific C-terminal antibody #C1557. As expected, Iba1⁺ microglia and GFAP⁺ astrocytes accumulated around amyloid plaques (Fig. 6G). Co-staining revealed a marked enrichment of Flower win–high astrocytes in the immediate plaque vicinity (Fig. 6H,I), with ∼45% of astrocytes near plaques exhibiting high Flower win expression compared with ∼7% in distal regions (Fig. 6J). In contrast, Flower win expression was largely absent from microglia, with only ∼4% of plaque-associated microglia and ∼0.5% of distal microglia displaying detectable Flower win levels (Fig. 6J).

Together, these data demonstrate a conserved, spatially restricted upregulation of Flower expression in astrocytes surrounding amyloid plaques in both human and mouse AD brains. The correlation between Flower expression and disease severity, combined with its selective enrichment in plaque-associated astrocytes, supports a model in which Flower-mediated fitness signaling contributes to astrocyte responses and tissue resilience in the neurodegenerative microenvironment.

### Flower win signaling enhances astrocytic Aβ uptake and protects neurons under amyloid stress

To determine whether Flower-mediated fitness signaling is functionally engaged during amyloid-induced neurodegeneration, we established an amyloidogenic neuron–glia co-culture model that recapitulates key aspects of Aβ accumulation and cytotoxicity. Primary cortical cultures were exposed to Aβ42 protofibrils, a toxic species generated under specific pre-incubation conditions (Condello *et al*, 2015; Sandberg *et al*, 2010).

Neuron–glia cultures were infected at day 1 in vitro (DIV1) with lentiviral constructs expressing either the Flower win isoform (mFwe2), the Flower C-terminal signaling domain, or mScarlet I as a control. On DIV7, cultures were treated with Alexa647-conjugated Aβ protofibrils (0.1 µM) and maintained for an additional six days before analysis (Fig. 7A). Aβ exposure induced marked cytotoxicity, with ∼45% of neurons and ∼35% of astrocytes becoming CC3-positive. Notably, overexpression of either mFwe2 or the C-terminal domain significantly reduced apoptosis in both cell types, restoring survival to near baseline levels comparable to untreated controls (Fig. 7B).

**Figure 7.**
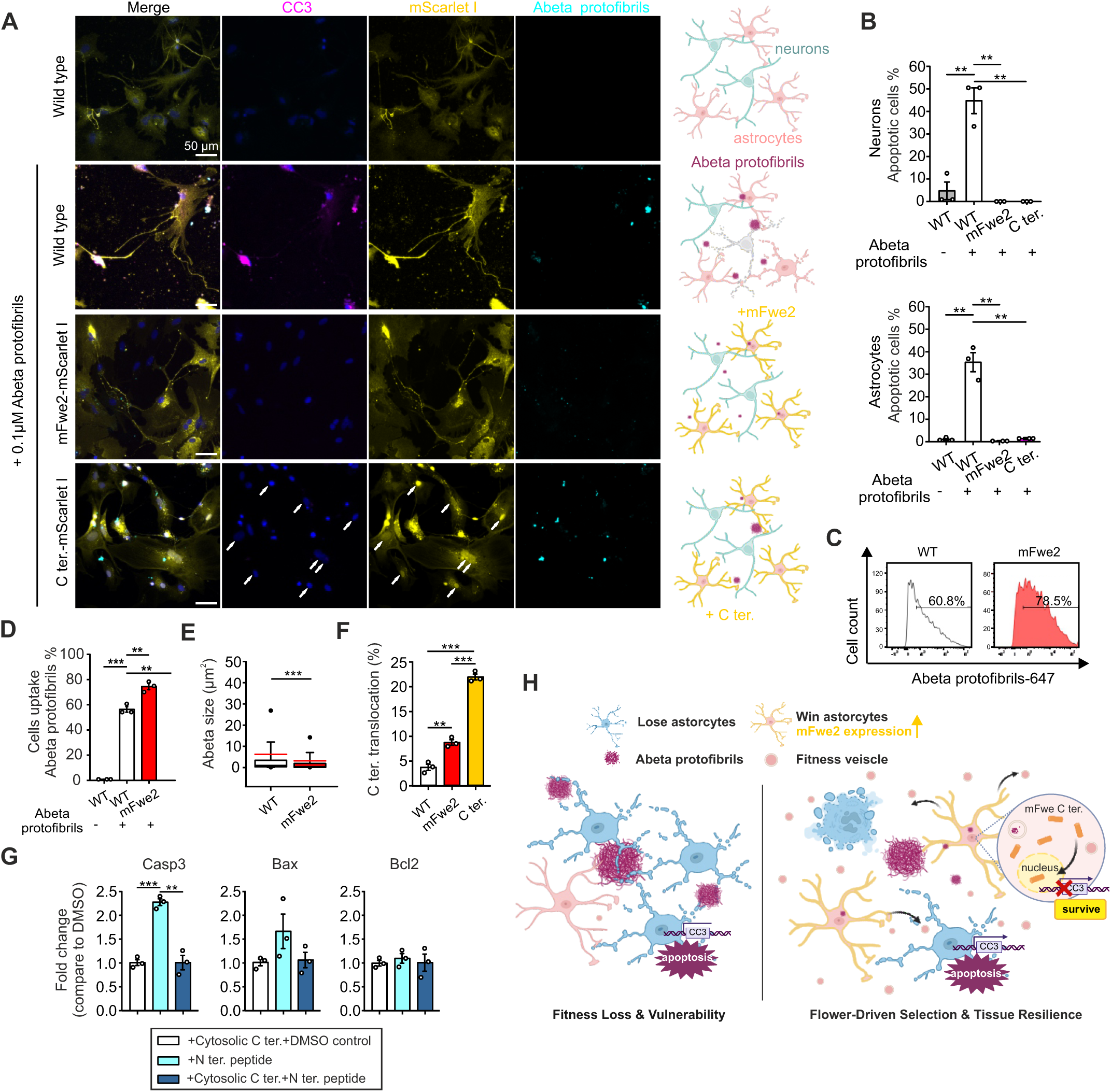
Flower “win” isoforms coordinate astrocyte fitness and Aβ clearance to promote neuroprotection. **(A)** Confocal images of transfected neuronal cultures treated with Aβ protofibrils-Alexa647. WT neuronal cultures were lentivirally infected at DIV1 with Flower-mScarlet I constructs (mFwe2-mScarlet I or C-terminal-mScarlet I) or a control mScarlet I vector (yellow). At DIV7, cultures were treated with or without Aβ protofibrils-Alexa647 (cyan). Neuronal cultures were fixed after 6 days and stained with CC3 antibody (magenta) to identify apoptotic cells. White arrows point to the translocated Flower C terminal protein. Scale bar, 50 µm. Right: schematic representation of the corresponding cultures and treatments shown on the left. **(B)** Quantification of CC3+ cell death from panel A. Data are presented as mean ± SEM. Statistical significance was determined using Student’s t-test: **p < 0.01. **(C)** Flow cytometric analysis of Aβ protofibril uptake by glial populations. Neuronal cultures were trypsinized and analyzed for the proportion of Aβ-positive cells. WT vector control (white) and mFwe2-expressing cells (red) are shown (N = 3). **(D)** Quantification of Aβ-positive cell populations from (**C**). Data are presented as mean ± SEM. Statistical significance was determined using Student’s t-test; **p < 0.01, ***p < 0.001. **(E)** Analysis of Aβ aggregate size in vector-transfected control (WT) and mFwe2-expressing cultures after 6 days. Data are presented as box-and-whisker plots. Statistical analysis was performed using one-way ANOVA followed by Dunn’s post hoc test; ***p < 0.001 (N = 3). **(F)** Quantification of Flower C-terminal nuclear translocation from confocal images shown in (**A**). **(G)** Quantitative RT–PCR analysis of apoptosis-related genes in cells overexpressing the cytosolic Flower C terminus under N-terminal peptide–induced death signaling. WT astrocytes were transfected with mFwe1 alone or co-transfected with the cytosolic C-terminal fragment to assess protective effects in unfit cells. Cells were treated with the N-terminal peptide, and expression of apoptosis-related genes (Caspase-3, Bax, and Bcl2) was analyzed after 24 h (N = 3). Data are presented as mean ± SEM. Statistical analysis was performed using two-way ANOVA followed by Holm–Šídák post hoc test; **p < 0.01, ***p < 0.001. **(H)** Model of Flower-mediated fitness surveillance and astrocyte-mediated neuroprotection. Schematic illustrating the bifurcated mechanism by which the Flower “win” isoform (mFwe2) coordinates astrocyte fitness and Aβ clearance. Left: In the absence of Flower-mediated selection, amyloid stress leads to widespread fitness loss and stochastic apoptosis, leaving the neural environment vulnerable. Right: Overexpression of mFwe2 reprograms the population: “win” astrocytes secrete Flower-positive EVs to eliminate distal, unfit “loser” cells while concurrently enhancing Aβ uptake. Internally, the C-terminal fragment translocates to the nucleus to repress Caspase-3 transcription, providing the cell-intrinsic survival necessary to maintain a protective astrocytic shield around plaques. Together, these pathways promote a robust state of tissue resilience in the AD brain.

Given the central role of astrocytes in Aβ uptake and clearance(Sollvander *et al*, 2016), we asked whether Flower signaling promotes a “clearance-competent” astrocyte state. Flow cytometry of surviving glia revealed that while approximately 60% of WT astrocytes internalized Aβ protofibrils, this proportion increased to ∼75% in astrocytes expressing mFwe2, indicating enhanced uptake capacity (Fig. 7C,D). Confocal imaging further demonstrated that mFwe2 expression significantly reduced the size of intracellular Aβ aggregates (Fig. 7E). Consistent with this specialized phenotype, mFwe2 induced the expression of astrocyte activation markers *(Gfap, Lcn2*), lipid transport regulators (*Abca1, Lrp1*), and engulfment receptors (*Megf10, Mertk*) (Fig. EV7A-E).

Notably, under amyloid stress, the C-terminal domain exhibited a pronounced intracellular redistribution, with significant nuclear enrichment observed in both mFwe2- and C-terminal-expressing cells (Fig. 7A,F). To test whether this intracellular pool directly suppresses apoptotic signaling, we challenged “unfit” (mFwe1-expressing) astrocytes with pro-apoptotic N-terminal peptides. Co-expression of the cytosolic C-terminal domain significantly attenuated the induction of *Caspase-3* mRNA, supporting an integrated neuroprotective model (Fig. 7G). In this framework, Flower “win” isoform-expressing astrocytes coordinate a bifurcated response: extracellularly, they enhance Aβ uptake while potentially secreting “fitness vesicles” (EVs) to eliminate distal “loser” cells; intracellularly, the C-terminal fragment translocates to the nucleus to repress the apoptotic transcriptional program. This dual regulation facilitates the selection of unfit cells while ensuring the survival of “win” cells around amyloid plaques, thereby establishing a robust astrocytic shield against pathology (Fig. 7H).

Together, these results demonstrate that Flower win signaling reprograms astrocytes toward a neuroprotective state, identifying the C-terminal domain as a key signaling module linking cellular fitness surveillance to tissue resilience during neurodegeneration.

## DISCUSSION

Our study uncovers a previously unrecognized mechanism by which astrocyte fitness is regulated through extracellular vesicle (EV)–mediated Flower signaling. While cellular fitness surveillance has traditionally been considered a contact-dependent process, our data demonstrate that Flower isoforms can function through non–cell-autonomous communication. Given the conserved role of fitness selection across development, aging, and disease(Coelho & Moreno, 2019; Levayer *et al*, 2016; Marques-Reis & Moreno, 2021; Merino *et al*., 2016; Merino *et al*., 2015; Moreno *et al*, 2015), this mechanism may represent a broader principle of tissue-level quality control that extends beyond neurodegeneration to tumor biology(Madan *et al*., 2019; Parker *et al*, 2021).

Mechanistically, Flower integrates opposing functional signals within a single transmembrane protein. The N-terminal region promotes the elimination of less-fit cells by activating intrinsic apoptotic pathways involving *Bax* and *Caspase-3* (Chipuk & Green, 2008; Kalkavan & Green, 2018), whereas the C-terminal domain antagonizes this death signaling. This intramolecular functional polarity distinguishes Flower from multifunctional regulatory proteins such as p53, Notch, or Bcl-2 family members (Delbridge *et al*, 2016; Kannan *et al*, 2013; Maddika *et al*, 2007; Vousden & Prives, 2009; Xie *et al*, 2014), which typically rely on post-translational modification, alternative splicing, or interaction networks to mediate opposing functions.

Our topology analyses further revise previous models proposing the constitutive extracellular orientation of Flower termini (Merino *et al*., 2013; Rhiner *et al*., 2010). Using pH-sensitive tagging, we demonstrate that Flower domains are predominantly cytoplasmic in intact cells but become extracellularly exposed specifically upon incorporation into vesicles. This suggests that intercellular communication occurs through EV-mediated molecular presentation. Notably, the identification of Flower-positive, CD63-negative vesicles defines a specialized population we term “fitness vesicles.” Ultrastructural and colocalization analyses suggest these vesicles originate from a non-canonical biogenesis pathway distinct from classical MVB-derived exosomes. While the precise molecular machinery governing the biogenesis and unconventional secretion route of Flower remains to be fully elucidated, our data suggest a pathway that is highly sensitive to cellular fitness states and independent of traditional endosomal sorting. Similar specialization has been reported in other systems, such as lamellar body–like organelles mediating cargo delivery in keratinocytes (Rudd *et al*, 2025), suggesting an evolutionary conservation of specialized signaling vesicles for tissue quality control. These observations align with emerging views that EVs function primarily in cellular homeostasis by exporting stress-associated cargo rather than serving as high-efficiency cytoplasmic delivery vehicles (Ngo *et al*, 2025).

Isoform-specific trafficking emerged as a key determinant of competitive outcome. Despite comparable intracellular expression, the win isoform mFwe2 was secreted approximately 25-fold more efficiently than mFwe1. This suggests a secretion-based “winner-take-all” dynamic, where “win” astrocytes actively “police” the tissue by broadcasting their signaling territory. Across our experimental paradigms, the elimination of less-fit astrocytes consistently reached ∼35–40%, likely reflecting intrinsic heterogeneity and a survival threshold within astrocyte networks.

The spatial enrichment of Flower in astrocytes surrounding Aβ plaques in both human AD brains and APP-transgenic mice (Sanchez-Mico *et al*., 2021) confirms the clinical relevance of this pathway. *CACFD1* expression increased progressively with Braak stage, suggesting localized activation of fitness surveillance within the neurodegenerative microenvironment. Functionally, *mFwe2* signaling reprograms astrocytes toward a neuroprotective clearance state, characterized by enhanced Aβ uptake and the induction of markers like *Gfap, Lcn2*, and engulfment receptors (*Megf10, Mertk*).

We propose that the functional coordination of Flower is governed by a “spatial paradox” resolved by secretion-dependent topology and stress-induced proteolysis. Our results demonstrate that while both termini are sequestered cytoplasmically in intact astrocytes, incorporation into “fitness vesicles” exposes them extracellularly—a transition critical for the C-terminal domain to act as a surface-masking antagonist that neutralizes N-terminal pro-apoptotic signals. Simultaneously, we find that the C-terminus translocates to the nucleus under amyloid stress to repress Caspase-3 (Fig. 7). We hypothesize that amyloid stress triggers a proteolytic cleavage event—potentially mediated by secretases or calpains—allowing the C-terminal fragment to pivot from its role as an extracellular “surface-shield” on EVs to an intracellular “nuclear-repressor” within the host cell. By coupling these spatially distinct mechanisms, Flower signaling ensures that “win” astrocytes are internally “immune” to the very death signals they broadcast to eliminate unfit neighbors.

The discovery that *CACFD1* expression is low in the healthy human brain but progressively upregulated in correlation with Braak stage severity reveals a fundamental principle of human brain resilience. We propose that Flower-mediated fitness surveillance serves as a “latent safeguard“—a homeostatic mechanism that remains largely quiescent during health but is specifically mobilized to gate tissue integrity under proteotoxic stress. By inducing a “win” phenotype in astrocytes surrounding plaques, the human brain attempts to construct a biological “firewall” that isolates pathology. The transition from incipient to severe Alzheimer’s Disease may thus be governed by the exhaustion of this fitness-selection machinery, where the astrocyte-led resilience can no longer keep pace with amyloid-induced fitness loss.

This modularity suggests a transformative “two-sided” therapeutic platform. The N-terminal “clearance” signal represents a tool for precision medicine, harnessing fitness-sensing machinery to force the competitive elimination of deleterious cells, such as tumors, by the healthy stroma. Conversely, the C-terminal “rescue” signal offers a blueprint for stabilizing fragile neurons. Delivering C-terminal peptides could provide a critical window of “transcriptional resilience,” providing reprogrammed glia the necessary time to clear the toxic microenvironment and restore neuronal functional integrity. This paradigm shifts the focus from merely slowing cell death to actively fostering a resilient cellular environment.

While this study focuses on astrocyte–astrocyte interactions, Flower-dependent signaling may also operate at neuron–glia interfaces. Given neurons’ sensitivity to proteotoxic stress (Condello *et al*., 2015; Sandberg *et al*., 2010), Flower-containing vesicles from astrocytes could modulate neuronal vulnerability by facilitating the removal of compromised cells or delivering pro-survival cues. Although our mechanistic conclusions rely on disease-relevant overexpression, the conserved upregulation in human AD tissue highlights Flower as a master regulator of glial resilience.

In summary, our findings establish Flower as a central regulator of astrocyte fitness and neuronal survival in Alzheimer’s disease. Through isoform-specific trafficking and EV-mediated communication, Flower orchestrates competitive survival decisions that shape glial responses to neurodegenerative stress. This non-canonical signaling paradigm positions Flower as a promising therapeutic target for enhancing glial support and preserving neuronal integrity.

## Methods

### Reagents and tools table

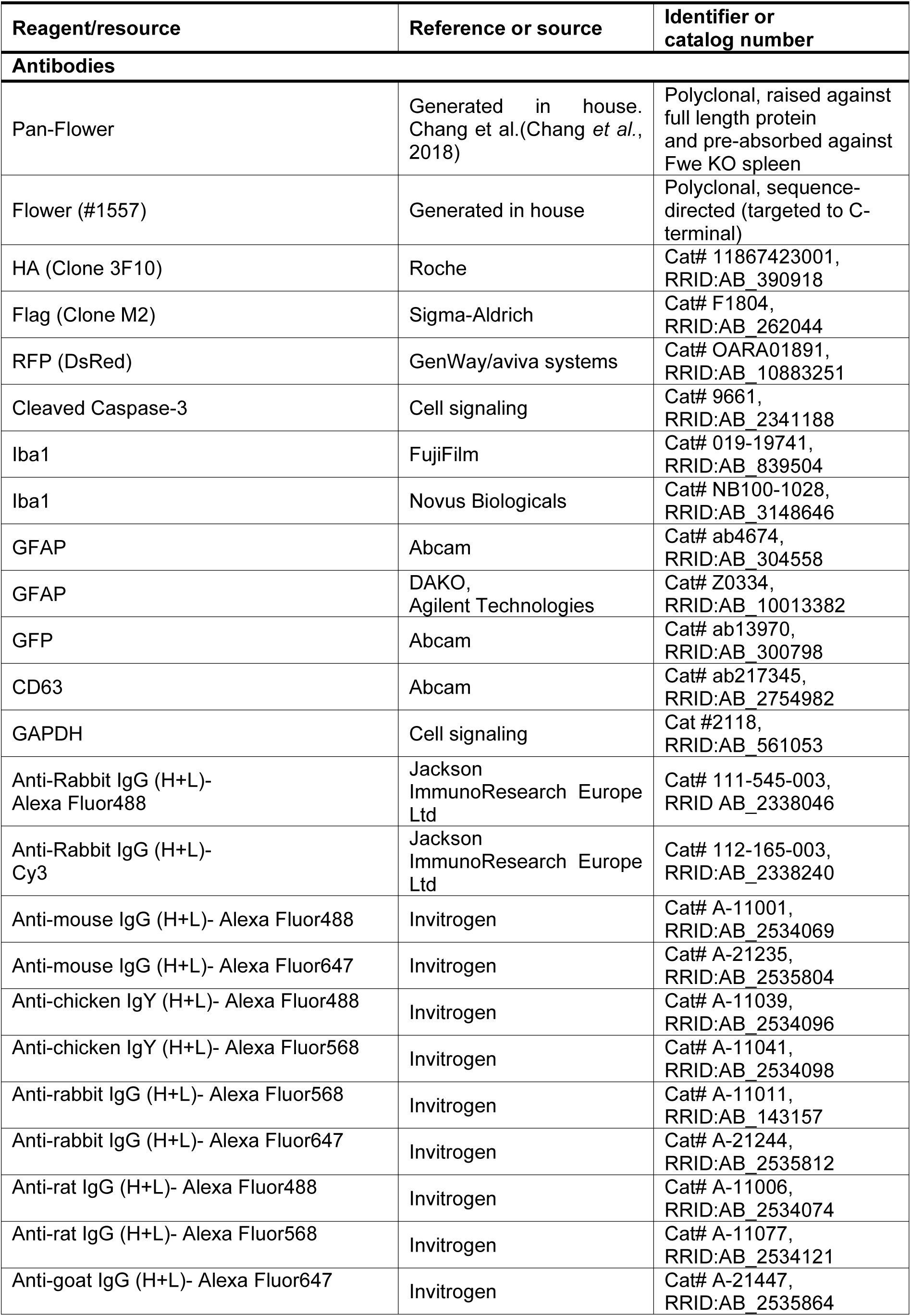
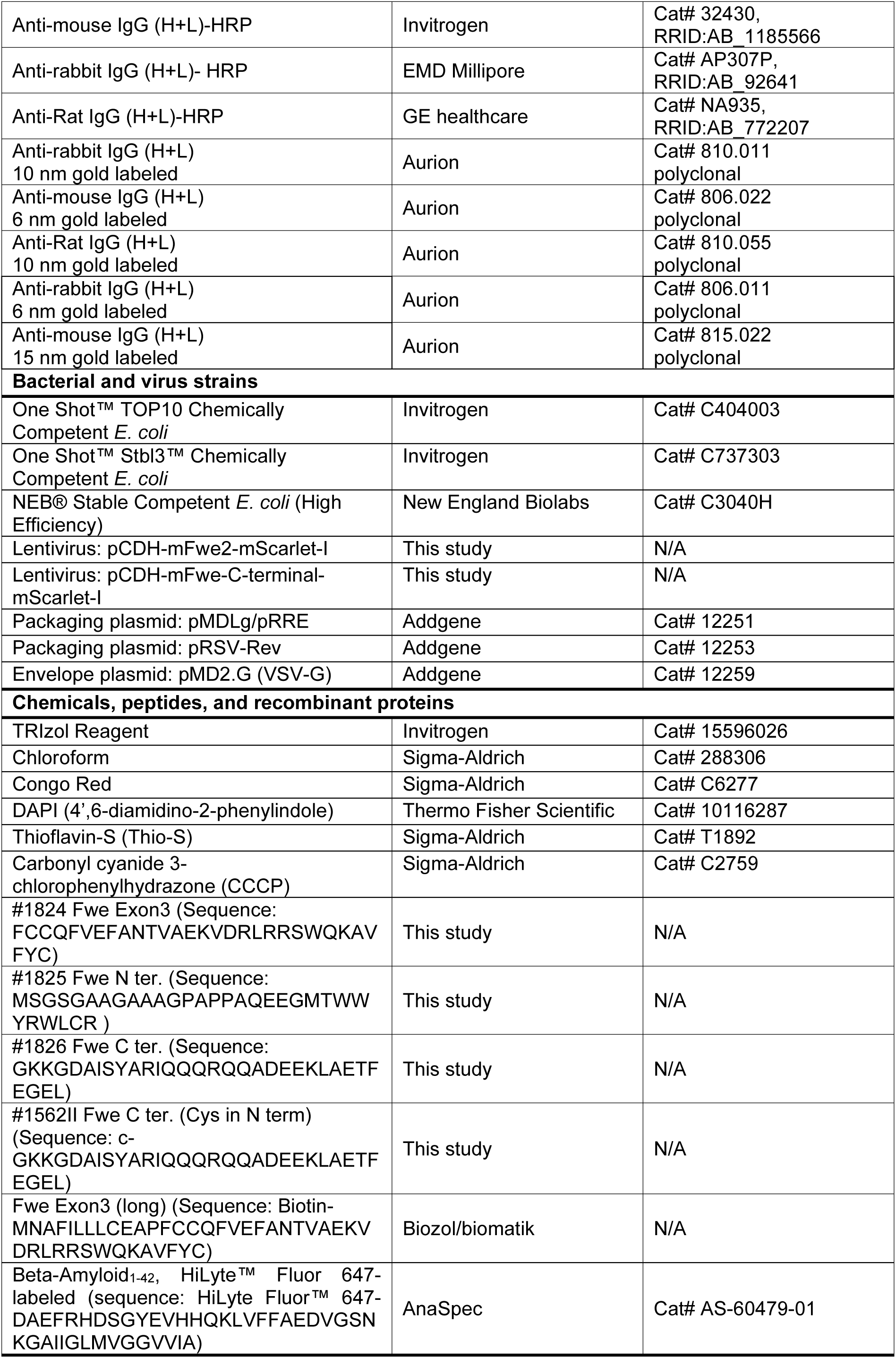
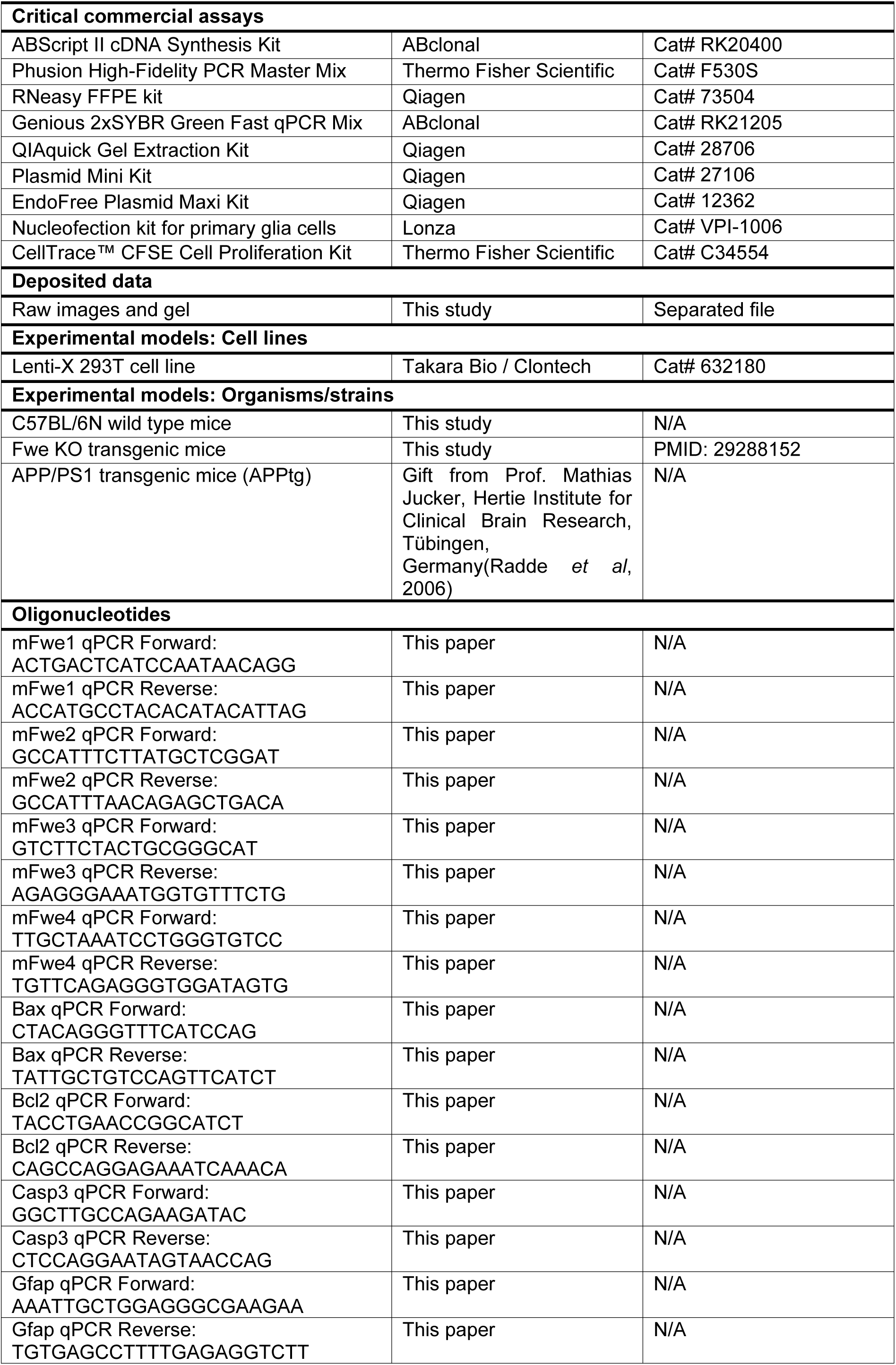
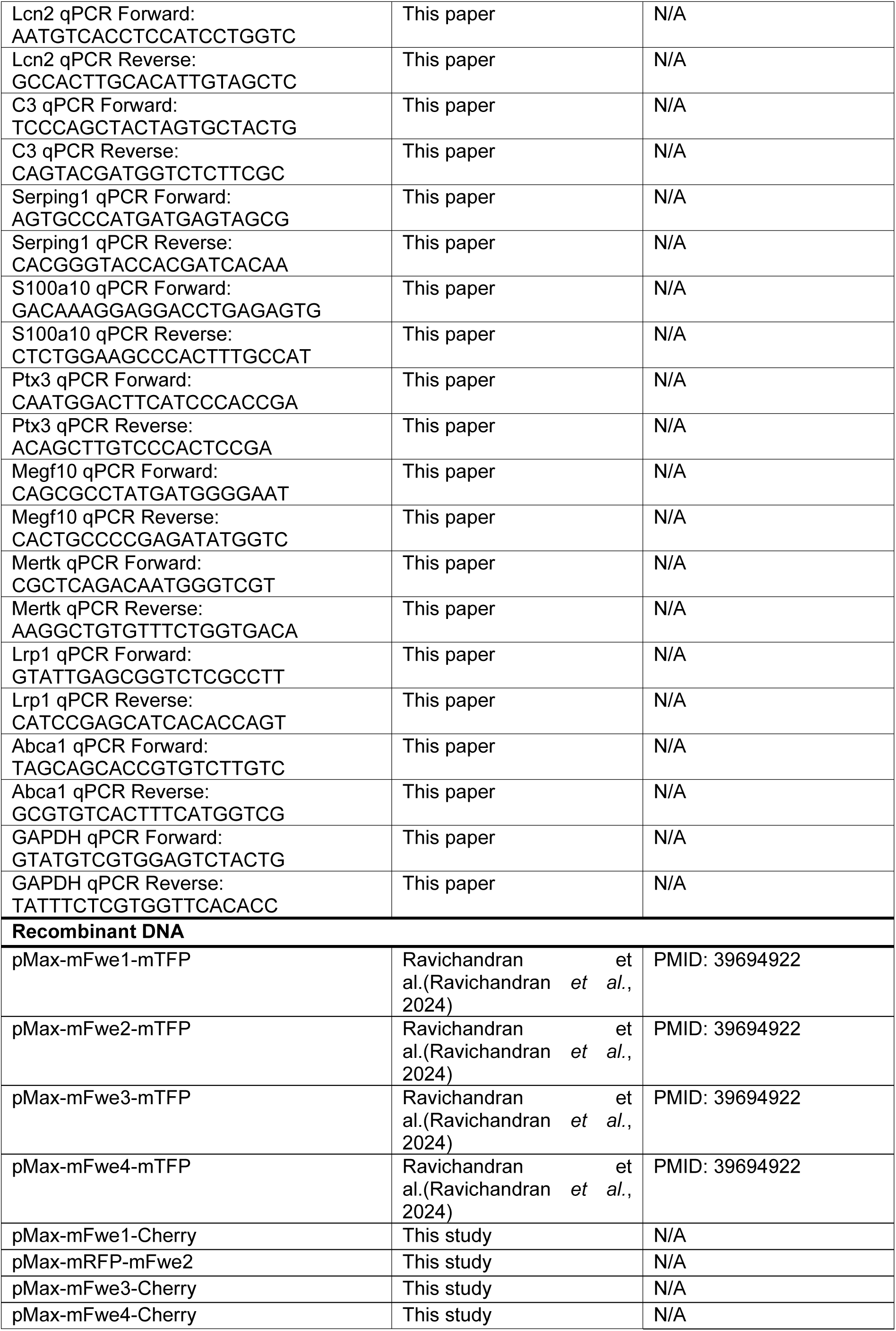
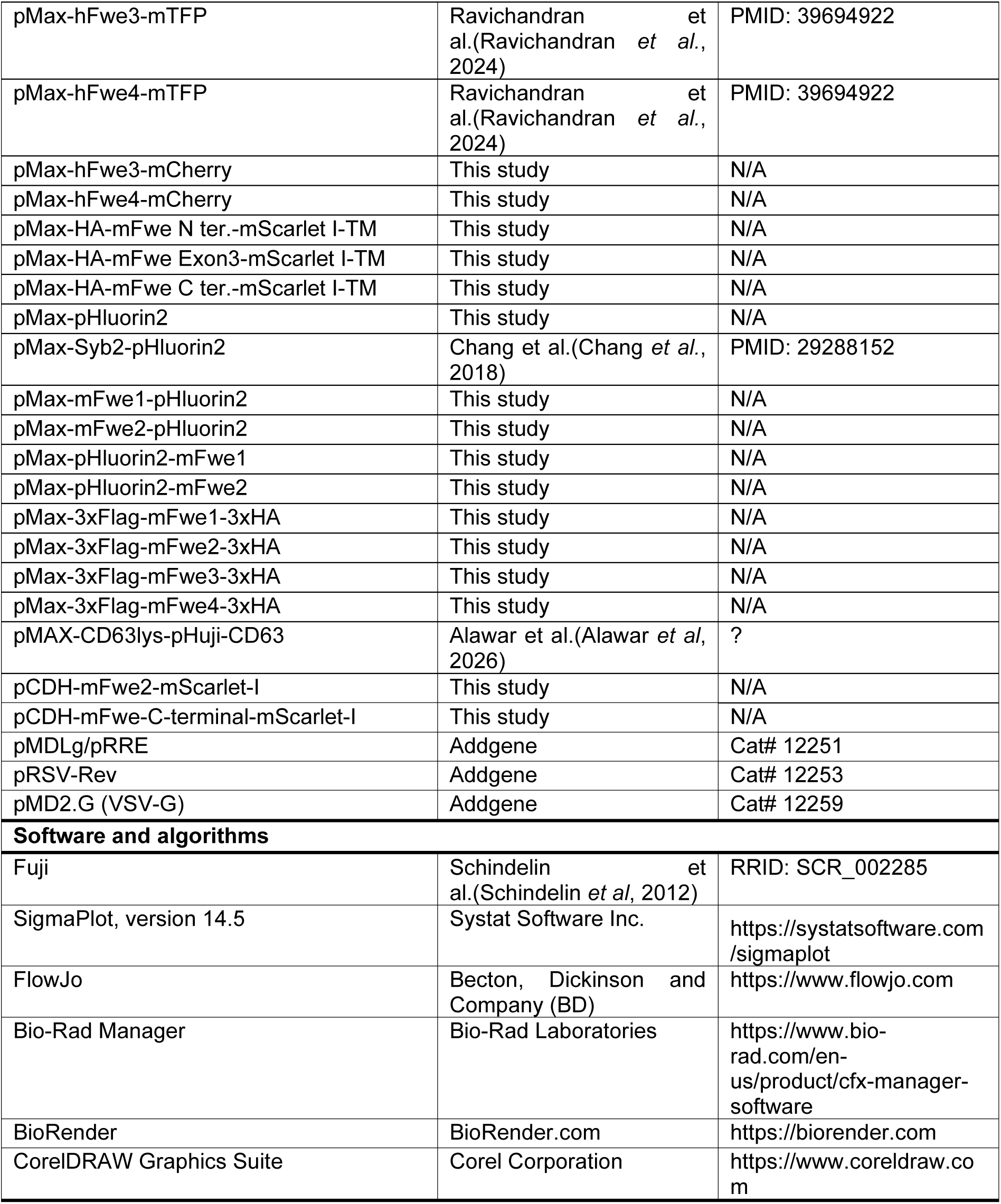

### Mice

Fwe KO mice were generated as previously described(Chang *et al*., 2018). APP/PS1 transgenic mice (APP^tg^), which overexpress human mutated APP (KM670/671NL) and PS1 (L166P) under the control of Thy1 promoters (PMC1559665), were kindly provided by M. Jucker (Hertie Institute for Clinical Brain Research, Tübingen, Germany). The KO, transgenic and WT mice used in this study were on a C57BL/6N background. Fwe KO pups used for astrocyte isolation were postnatal day 0–2 (P0–P2). Adult WT and APP mice used for brain tissue staining and RNA analysis were 60-70 weeks old and included both sexes. Animals were housed at 22 °C with 50–60% humidity under a 12 h light/dark cycle. All experimental procedures were approved by the local regulatory authority of Saarland (Landesamt für Verbraucherschutz; protocols AZ: 2.4.1.1/11-2021 to E.K. and VM2023-05 to Y.L.) and carried out in accordance with institutional and governmental guidelines.

### Human postmortem brain tissue

Formalin-fixed, paraffin-embedded hippocampal sections from autopsy-confirmed Alzheimer’s disease (AD) patients were obtained from the Department of Neuropathology, Saarland University, Germany. The use of human tissue was approved by the Ärztekammer des Saarlandes (Ethics Committee of the Medical Association of Saarland), approval number 106/25. These paraffin-embedded hippocampal sections were used for immunostaining experiments as described in the Method Details section.

### Primary cortical neuron and astrocyte culture

Cortical neurons and astrocytes were isolated from P0–P2 mouse pups. Cerebral hemispheres were dissected in ice-cold Earle’s Balanced Salt Solution (EBSS; Gibco). Following removal of the meninges, cortices were minced into small fragments and enzymatically digested with 35 U/mL papain (Worthington) at 37 °C for 45 min, followed by gentle mechanical trituration to obtain a single-cell suspension. For neuronal culture, dissociated cells were seeded onto 25 mm glass coverslips placed in 6-well culture plates at a density of 3 × 10^5^ cells per coverslip. Coverslips were pre-coated with a solution containing 17 mM acetic acid, collagen I (Gibco), and poly-D-lysine (Sigma). On day in DIV 1, the culture medium was replaced to remove residual debris. Neurons were maintained in Neurobasal-A (NBA) medium supplemented with 10% FCS, 1% penicillin-streptomycin (Invitrogen), 1% GlutaMAX, and 2% B27 supplement (Gibco). Neurons were cultured for 9–12 days prior to experimental use. For astrocyte culture, dissociated cortical cells were seeded into T-75 culture flasks (one brain per flask), pre-coated with poly-L-ornithine (Sigma). On DIV1, flasks were rinsed and vigorously agitated with DPBS to remove debris and non-adherent cells. The remaining adherent astrocytes were cultured in Dulbecco’s Modified Eagle Medium (DMEM; Gibco) supplemented with 10% FCS and 1% penicillin-streptomycin for 9–14 days before use in experiments.

### Flower antibody generation

Details of Flower antibody generation have been previously described(Chang *et al*., 2018). In this study, two custom-made polyclonal antibodies were used. The first, referred to as Flower antibody #1557, was sequence-directed and raised against the C-terminal 18 amino acids shared by mFwe2 and mFwe4. The second, a pan-Flower antibody, was generated against the untagged full-length recombinant mFwe2 protein(Chang *et al*., 2018) and recognizes multiple epitopes across the Flower protein, including both N- and C-terminal regions.

### Peptides synthesis

Peptides were synthesized with an automated ResPepSL synthesiser (Intavis) using amide Rink resin as the solid phase and Fmoc-protected amino acids (Carbolution) for coupling according to the method of Merrifield(Merrifield, 1963). The resin-coupled peptides were cleaved and deprotected with trifluoroacetic acid (Sigma Aldrich) followed by precipitation with tert-butyl methyl ether (Fisher Scientific) and further analysis and purification by preparative HPLC (Merck) (purtity > 90%). Lyophilized peptides (Lyovac GT2, Finn-Aqua) were stored at 4 °C. Stock solutions of the peptides were prepared fresh and used the same day.

### Mouse brain slice preparation and staining

APP-transgenic mice and age-matched WT littermates were sacrificed by CO_2_ inhalation followed by decapitation. Brains were extracted and post-fixed in 4% paraformaldehyde (PFA) at 4 °C for 24 h. Fixed tissues were sectioned into 30 μm slices using a vibratome (Leica VT1000 S). Sections were washed three times with Dulbecco’s PBS (D-PBS), permeabilized with 0.1% Triton X-100 for 10 min, and blocked with 10% fetal calf serum (FCS) for 30 min at room temperature (RT). Primary antibodies rabbit anti-Iba1 (1:200, 019-19741; FujiFilm), goat anti-Iba1 (1:200, NB100-1028; Novus Biologicals), chicken anti-GFAP (1:1000, ab4674; Abcam) and polyclonal rabbit anti-Flower #1557 (1:200, homemade) were applied overnight at 4 °C, followed by incubation with corresponding Alexa Fluor-conjugated secondary antibodies (Invitrogen, 1:1000). Nuclei were counterstained with DAPI, and slices were mounted and imaged by confocal microscopy.

### Immunostaining of postmortem human brain sections from AD patients

Formalin-fixed, paraffin-embedded hippocampal sections from autopsy-confirmed Alzheimer’s disease (AD) patients were obtained from the Department of Neuropathology, Saarland University, Germany. The use of human tissue was approved by the Ärztekammer des Saarlandes (Ethics Committee of the Medical Association of Saarland), approval number 106/25. Following deparaffinization, slides were subjected to antigen retrieval in citrate buffer (pH 6.0) for 30 min. Endogenous peroxidase activity was quenched with 3% H₂O₂ in PBS, and sections were blocked for 1 h in 0.2% casein.

Immunostaining was performed using a rabbit polyclonal anti-GFAP antibody (1:500, Z0334; DAKO, Agilent Technologies, Waldbronn, Germany), followed by detection with an alkaline phosphatase (AP)-conjugated goat anti-rabbit IgG secondary antibody (DAKO) and visualization using the Vector Red Substrate Kit (SK-5100; Vector Laboratories, Newark, USA).

To assess astrocytic localization of Flower protein, serial sections were stained with a rabbit polyclonal anti-Flower antibody, followed by GFAP staining. Detection was performed with either AP- or horseradish peroxidase (HRP)-conjugated goat anti-rabbit IgG secondary antibodies (both from DAKO), and developed with either 3,3’-diaminobenzidine tetrahydrochloride (DAB; Sigma) or the Vector® Blue Substrate Kit (SK-5300; Vector Laboratories). Slides were counterstained with hematoxylin.

For fluorescence-based colocalization analysis, the same staining protocol was repeated using Alexa Fluor 488- or Cy3-conjugated goat anti-rabbit IgG secondary antibodies (Jackson ImmunoResearch Europe Ltd, Cambridgeshire, UK). Colocalization between Flower and GFAP was evaluated by confocal microscopy (LSM 780, Zeiss).

### Human Alzheimer’s disease transcriptome analysis

Transcriptome data from hippocampal specimens derived from patients with Alzheimer’s disease (AD) of varying disease severity, as well as from non-demented control individuals, were obtained from the Gene Expression Omnibus (GEO) database (Blalock *et al*., 2004) (accession number GSE1297). Expression values of CACFD1 were extracted from the dataset and normalized to z-scores by subtracting the dataset mean and dividing by the standard deviation. Differences in CACFD1 expression among groups were assessed using a two-tailed Kruskal–Wallis test. Statistical analyses were performed using GraphPad Prism version 10.2.3. The analysis protocol was approved by the Institutional Research Ethics Committee of China Medical University Hospital, Taiwan (CMUH113-REC1-167).

### Cell fitness assay

To assess the fitness effects of Flower isoforms, Flower KO astrocytes cultured for 9–12 days were used. Cells were detached using Trypsin-EDTA, washed three times with DPBS, and resuspended in 100 µL nucleofection solution (Basic Nucleofector Kit for Primary Mammalian Glial Cells, Lonza). Each nucleofection was performed with 4 × 10^6^ astrocytes and 5 µg plasmid DNA encoding fluorescently tagged win or lose Flower isoforms. Electroporation was carried out using a electroporation device (Lonza NucleofectorI/II/2b System). Following electroporation, equal numbers of the two transfected astrocyte populations were mixed and co-seeded into 6-well plates containing 25 mm coverslips (around 4 × 10^5^ mixed cells per coverslip) in the following combinations: win+win, win+lose, and lose+lose. To evaluate the role of specific Flower protein domains in cell competition, three additional constructs were generated, each encoding a truncated version of Flower fused to mScarlet I-TM, targeting either the N-terminus, Exon 3, or C-terminus. These constructs were individually transfected into Flower KO astrocytes and co-cultured with an equal number of Flower KO astrocytes expressing the lose isoform. After 24 h of co-culture, cells were fixed in 4% PFA for 10 min at RT. DAPI was added prior to fixation to label late apoptotic cells. Immunocytochemistry was performed using a rabbit anti-Cleaved Caspase-3 antibody (1:500, #9661; Cell Signaling) to detect early apoptotic cells.

For peptide-based assays, synthetic peptides corresponding to the N-terminus, Exon 3, and C-terminus regions of Flower were added to astrocyte cultures at a concentration of 1 µg/ml. Astrocytes expressing either win or lose isoforms were treated for 24 h prior to fixation and analysis. Finally, Live or fixed cells were imaged using a confocal microscope (LSM780, Zeiss) equipped with a 40×/1.3 NA objective. Cell counts and cell death analysis were performed manually using Fiji (ImageJ).

To evaluate the role of secreted Flower in cell fitness, supernatants from T-75 flasks were collected, and EVs were isolated by ultracentrifugation (see EV isolation). Approximately 20 µg of protein from the pelleted supernatant can be obtained from one flask (10 ml). Ten micrograms of pelleted EVs were applied to approximately 1 × 10^5^ unfit astrocytes expressing mFwe1-mCherry, which were seeded on a 12.5 mm coverslip for the fitness assay. After 24 h, cells were fixed and stained for CC3 to quantify apoptotic cells.

### Amyloid Beta1–42 protofibril generation and application

Synthetic Aβ_1-42_ peptides labeled with HiLyte Fluor 647 (Anaspec Inc.) were dissolved in 1% NH_4_OH and subsequently diluted in 1x PBS to a final concentration of 100 µM. The solution was incubated at 37 °C for 4 h to allow for protofibril formation. Protofibrils were then diluted to a final concentration of 0.1 µM and applied to cortical neuron cultures on DIV 7 for 6 days to assess cytotoxicity. WT cortical neuron cultures were transduced with lentiviruses encoding mFwe2-mScarlet I, mFwe C-terminal-mScarlet I, or cytosolic mScarlet I at DIV 1. Cells were then either fixed and immunostained for apoptosis analysis, collected for flow cytometry to assess Aβ_1-42_ protofibril uptake, or harvested for RNA extraction followed by qPCR analysis.

### Plasmid cloning

Plasmids encoding mouse Fwe1, Fwe2, Fwe3, and Fwe4, as well as human Fwe3 and Fwe4 with C-terminal mTFP tags, were generated in a previous study(Ravichandran *et al*., 2024). These constructs served as the backbone for the generation of additional tagged and truncated Flower expression plasmids used in this study.

To generate C-terminal mCherry fusion constructs (pMax-mFwe1-Cherry, pMax-mFwe3-Cherry, pMax-mFwe4-Cherry, pMax-hFwe3-mCherry, and pMax-hFwe4-mCherry), the mTFP sequence in the parental pMax-mFwe-mTFP plasmids was replaced with mCherry using XbaI and XhoI restriction sites. For pMax-mRFP-mFwe2, the mRFP sequence was cloned upstream of the mFwe2 coding sequence.

For generation of truncated Flower constructs fused to a transmembrane (TM) domain, the pDisplay-pHuji vector (#61556, Addgene), containing the PDGFR TM domain, was used as a template. The mScarlet-I sequence and PDGFR TM domain were subcloned into the pMax backbone to generate a pMax-mScarlet-I-TM vector. Truncated Flower sequences corresponding to the N-terminal region, exon 3, or C-terminal region were PCR-amplified using gene-specific primers. The C-terminal Flower fragment flanked by BglII and BamHI sites was synthesized (BioCAT) in pBluescript II SK(+). These fragments were subcloned into pMax-mScarlet-I-TM to generate pMax-HA-mFwe-N-ter-mScarlet-I-TM, pMax-HA-mFwe-Exon3-mScarlet-I-TM, and pMax-HA-mFwe-C-ter-mScarlet-I-TM constructs.

To generate dual-tagged Flower constructs, synthetic DNA fragments encoding either a 3×Flag tag (flanked by ClaI and EcoRI sites) or a 3×HA tag (flanked by XbaI and XhoI sites) were obtained from BioCAT. The 3×Flag sequence replaced mScarlet-I, and the 3×HA sequence replaced mTFP in the corresponding pMax-mFwe plasmids, resulting in pMax-3×Flag-mFwe1-3×HA, pMax-3×Flag-mFwe2-3×HA, pMax-3×Flag-mFwe3-3×HA, and pMax-3×Flag-mFwe4-3×HA.

The pmax-CD63-pHuji was generated by subcloning from pCMV-CD63-pHuji into pMax with the restriction sites EcoRI and XbaI. Its size was 4.282 kb. pCMV-CD63-pHuji was a generous gift from Frederik Verweij (Centre de Psychiatrie et neurosciences, Amsterdam/Paris). In the construct the pH sensitive fluorescent marker protein was located to the first extracellular loop of the tetraspanins facing the acidic lumen of the organelles(Verweij *et al*., 2018).

For pHluorin2-based constructs, pHluorin2 was PCR-amplified and cloned either C-terminally or N-terminally. For C-terminal fusions (pMax-mFwe1-pHluorin2 and pMax-mFwe2-pHluorin2), the mTFP sequence was replaced by pHluorin2 using XbaI and XhoI. For N-terminal fusions (pMax-pHluorin2-mFwe1 and pMax-pHluorin2-mFwe2), the Halo tag in pMax-Halo-mFwe constructs was replaced with pHluorin2 using EcoRI and NheI. pMax-Syb2-pHluorin2 was generated previously⁴.

For lentiviral expression, the pCDH-EF1 vector (#72266, Addgene) was used. mFwe2 was cloned into pCDH-EF1 using EcoRI and XbaI, followed by insertion of mScarlet-I to generate pCDH-mFwe2-mScarlet-I. To generate pCDH-mFwe-C-terminal-mScarlet-I, the C-terminal Flower fragment was PCR-amplified and used to replace the full-length mFwe2 sequence in the pCDH-mFwe2-mScarlet-I backbone. All constructs were verified by Sanger sequencing using gene-specific primers (Microsynth Seqlab).

### Surface plasmon resonance binding analysis

Series Sensor Chips CM5 were obtained from Cytiva (Freiburg, Germany). Amine coupling reagents N-(3-dimethylaminopropyl)-N’-ethylcarbodiimide hydrochloride (EDC) and N-hydroxysuccinimide (NHS), along with ethanolamine hydrochloride, were purchased from Merck KGaA (Darmstadt, Germany). HEPES buffer, NaCl, and NaOH were sourced from Carl Roth (Karlsruhe, Germany), Grüssing GmbH (Filsum, Germany), and VWR Chemicals (Darmstadt, Germany), respectively. TWEEN 20 and dimethyl sulfoxide (DMSO) were acquired from Sigma-Aldrich (Steinheim, Germany) and Thermo Fisher Scientific (Dreieich, Germany). For SPR analysis, HBS-P (0.01 M HEPES, 0.15 M NaCl, 0.05% TWEEN 20) served as running buffer during surface preparation. All glassware was rinsed with 50 mM NaOH and Milli-Q water before use. Ligand 1562II was immobilized onto a CM5 sensor chip via amine coupling using a BIACORE T200 system (Cytiva, Freiburg, Germany). Carboxyl groups were activated with a 1:1 mixture of 391.2 mM EDC and 100 mM NHS (10 μl/min, 7 min). 1562II (200 µg/ml in 10 mM HEPES, pH 5.5) was injected for 2 min (10 μl/min), achieving an immobilization level of about 4690 RU. Residual esters were blocked with 1 M ethanolamine-HCl (pH 8.5, 7 min). A reference surface was prepared by activation and immediate deactivation without ligand.

Binding of 1825 and 1826 to immobilized 1562II was assessed at 25 °C using HBS-P with 3 % DMSO as running buffer. Solvent correction was performed using DMSO dilutions (2.5 - 3.8 %). Analytes were prepared at 15, 30, and 40 µM in running buffer and injected in triplicate. Each cycle included a 180 s association and 300 s dissociation phase at 30 μl/min. Surfaces were washed with running buffer and cleaned with 50 % DMSO. Regeneration was achieved using 5 M NaCl followed by deionized water. Data were processed with BIACORE T200 Evaluation Software v3.2.1, applying double referencing and solvent correction prior to curve fitting.

### Harvest of secreted Flower fractions and extracellular vesicles

WT astrocytes cultured for 9–12 days were electroporated with constructs expressing mFwe2-pHluorin2, 3×Flag-mFwe1-3×HA, or 3×Flag-mFwe2-3×HA. Following electroporation, 4 × 10^6^ cells were cultured in a T-75 flask containing 10 ml of culture medium for 2–3 days. The medium was supplemented with 10% exosome-depleted FCS, prepared by ultracentrifuging heat-inactivated FCS (Gibco) at 100,000 × g for 16 h at 4 °C to remove serum-derived EVs. To harvest astrocyte-derived EVs, conditioned medium was collected and centrifuged at 300 × g for 10 minutes to remove cells and large debris, followed by an additional centrifugation step at 2,000 × g for 20 min to further eliminate residual cell debris and dead cells. The supernatant was then filtered through a 0.22 μm syringe filter (Millipore) to eliminate residual debris and large vesicles. Sequential ultracentrifugation was performed: first at 10,000 × g for 40 minutes at 4 °C to remove apoptotic bodies, followed by 100,000 × g for 1 h and 15 minutes at 4 °C to pellet secreted Flower-containing fractions and EVs. The resulting pellet was resuspended in D-PBS and washed by a second ultracentrifugation step at 100,000 × g for 1 h and 15 min at 4 °C to reduce contaminating soluble proteins. All ultracentrifugation steps were carried out using Ultra-Clear centrifuge tubes (Beckman Coulter) in an Optima XPN-80 ultracentrifuge (Beckman Coulter). For analysis of secreted Flower and EV content, pellets were resuspended in D-PBS and applied to 12.5 mm coverslips pre-coated with poly-L-ornithine. After a 1 h incubation at RT, the supernatant was carefully removed to avoid disturbing the settled EVs. The samples were then fixed with 4% PFA for 20 min, followed by immunostaining.

### EV particle size analysis

EVs harvested from GFP, mFwe1 or mFwe2-overexpressing WT astrocyte cultures were analyzed by asymmetric-flow field-flow fractionation (AF4) using an Eclipse system (Wyatt Technology, Santa Barbara, CA, USA) coupled to an Agilent 1260 Infinity II LC (Agilent Technologies, Waldbronn, Germany). Online detection included multi-angle light scattering (MALS; DAWN, 18 angles, 658 nm, Wyatt Technology), differential refractive index (dRI; Optilab, Wyatt Technology), and UV–Vis (Agilent 1260 DAD) measurements. Instrument control and sequence execution were performed using VISION v2.x (Wyatt Technology), and data were processed in ASTRA v7.3 (Wyatt Technology). Following a focus step, samples were eluted under programmed cross-flow at a constant detector flow. Geometric radii were calculated slice-wise in ASTRA using a spherical Lorenz–Mie model, specifying the material refractive index, and entering the refractive index increment (dn/dc) or extinction coefficients as appropriate for dRI or UV–Vis detection. Absolute particle number concentrations were obtained using ASTRA’s “Number from Light Scattering” analysis, with number density per slice integrated over each elution peak and normalized to the injected volume. Prior to sample analysis, detectors were aligned and normalized according to manufacturer procedures, membranes were conditioned, and a monodisperse polystyrene latex standard was used to verify retention behavior and MALS response. Mobile-phase blanks were injected between samples to monitor carryover and baseline stability(Cho & Hackley, 2010).

### Immunogold labeling and electron microscopy

After size analysis and quantification, 40 µl of each EV sample were fixed with PFA (2% final concentration). For immunogold analysis, 7 µl of the fixed EVs were dropped onto pioloform-coated 200-mesh copper grids for 45 min. After several washing steps with D-PBS and 50 mM glycine in D-PBS for 10 min, the grids were incubated in blocking solution (Aurion) for another 10 min. The primary antibodies anti-HA (1:10), anti-Flag (1:10), anti-RFP (1:50) and the pre-absorbed anti-Flower (1:2) were diluted in incubation solution (D-PBS supplemented with 0.1 % BSAc (Aurion), pH 7.4), and the grids were incubated for 2 hours at 20 ±2 °C. After several washes the grids were incubated for 1 hour at 20 ±2 °C with the secondary gold-conjugated antibodies goat anti-rabbit (6 nm or 10 nm), goat-anti rat (10 nm), goat anti-mouse (6 nm or 15 nm) (Aurion), diluted in incubation solution (1:30). The samples were then washed in several drops of D-PBS, fixed in 1% glutaraldehyde, rinsed with six drops of destilled water, and contrasted with uranyl acetate. The grids were analyzed with a Tecnai G2 Biotwin electron microscope (Thermo Fisher Scientific) and electron micrographs were acquired using Olympus iTEM 5.0 image software (build1243).

### Western blot analysis

Cells were lysed on ice using a buffer containing 50 mM Tris-HCl (pH 7.4), 1 mM EDTA, 250 µM PMSF, 1% Triton X-100, 150 mM NaCl, 1 mM DTT, 1 mM sodium deoxycholate, and a protease inhibitor cocktail (Roche). Cell lysis was facilitated by mechanical scraping, followed by incubation on ice for 30 min. Lysates were centrifuged at 16,000 × g for 10 min at 4 °C, and the supernatants were collected for subsequent analysis. Protein concentrations were determined using the Pierce 660 nm Protein Assay (Thermo Fisher Scientific) according to the manufacturer’s instructions.

Equal amounts of protein (30 µg per lane for cell lysates, 10 µg per lane for secreted Flower fractions from conditional medium) were resolved by SDS–PAGE using NuPAGE precast gels (Invitrogen) and transferred onto nitrocellulose membranes. Membranes were blocked for 30 min at RT in TBS-T buffer (20 mM Tris-HCl, 150 mM NaCl, 0.05% Tween-20, pH 7.4) containing 5% (w/v) non-fat dry milk. Membranes were then incubated with primary antibodies rabbit polyclonal anti-Pan Flower (1:1000, homemade), rat anti-HA (1:1000, clone 3F10, Roche), mouse anti-Flag (1:1000, clone M2, Sigma-Aldrich), rabbit anti-GAPDH (1:5000, clone 14C10, Cell Signaling) and rabbit anti-CD63 (1:500, clone EPR21151, Abcam) overnight at 4 °C. After washing three times in TBS-T, membranes were incubated with HRP-conjugated secondary antibodies for 45 min at RT. Signal detection was performed using enhanced chemiluminescence reagents (SuperSignal West Dura Chemiluminescent Substrate, Thermo Fisher Scientific), and images were acquired using the FluorChem E documentation system (BioLabTec GbR).

### Immunofluorescence

Cells grown on coverslips were fixed in 4% PFA for 10 min at RT and subsequently washed three times with D-PBS. Permeabilization was performed in D-PBS containing 0.1% Triton X-100 (Roth) for 10 min. Samples were then incubated in blocking buffer (D-PBS supplemented with 2% BSA) for 30 min to reduce non-specific binding. Primary antibodies rabbit polyclonal anti-Pan Flower (1:100, homemade), rat anti-HA (1:100, clone 3F10, Roche), and mouse anti-Flag (1:100, clone M2, Sigma-Aldrich), rabbit anti-RFP (DsRed) (1:200, OARA01891, GenWay/aviva systems) were diluted in blocking buffer and applied for 1 h at RT. Following three washes with D-PBS, samples were incubated with the appropriate Alexa Fluor-conjugated secondary antibodies (1:1000, Invitrogen) for 45 min at RT in the dark. After three additional washes with D-PBS, samples were counterstained with or without DAPI (1:1000) and mounted using Fluoromount-G Mounting Medium (Invitrogen). Cell death was analysed using CC3 and DAPI markers by confocal microscopy with a 40×/1.3 NA objective. Maximum-intensity projections were used to illustrate Flower protein localization and cell death in the cultures.

Secreted Flower particles were analysed by super-resolution SIM using a 63×/1.4 NA objective. SIM imaging were performed on a Zeiss Elyra PS.1 microscopic SIM System with Zen 2010 software for device control and high-resolution image processing. Images were acquired as z-stacks with 0.15-µm intervals through the entire particles. Single-plane images were used to illustrate Flower protein localization on the particles.

### Amyloid plaques extraction by laser capture dissection

Freshly isolated brains from APP-transgenic mice were fixed in 4% PFA overnight and subsequently processed for paraffin embedding. Fixed tissues were dehydrated through a graded ethanol series (70%, 80%, and 95% ethanol for 1.5 h each), followed by three washes in 100% ethanol (2 h each). Samples were cleared in xylene (three times for 1.5 h), then infiltrated with paraffin wax at 60 °C (two rounds of 1.5 h each). Embedded tissues were oriented in molds and cooled at room temperature to solidify.

Paraffin blocks were sectioned at 5 μm thickness using a microtome. Sections were floated on a 63–65 °C water bath to remove wrinkles, then mounted onto RNase-free PEN membrane-coated glass slides (VWR, Cat. No. 76414-898) for laser microdissection (Leica). Slides were air-dried and stained with Congo Red to label amyloid plaques and Hematoxylin to visualize cellular architecture prior to RNA extraction.

Laser capture microdissection was performed using a Leica LMD system. Congo Red–positive amyloid plaques were identified under fluorescence illumination and precisely cut using the UV laser. Microdissected tissue fragments were collected directly into RNase-free microcentrifuge tube caps containing lysis buffer for RNA isolation. To minimize RNA degradation, dissections were completed within 1–2 h per slide, and slides were kept dry and protected from light. At least 100 ng total RNA was collected per sample.

RNA was extracted from microdissected tissues using the RNeasy FFPE Kit (QIAGEN) following the manufacturer’s instructions. Briefly, tissues were lysed in PKD buffer, treated with Proteinase K, incubated at 56 °C for 15 min followed by 80 °C for 15 min, and cooled on ice. Samples were centrifuged at 20,000 × g for 15 min, and the supernatant was treated with DNase I. RNA was precipitated, bound to an RNeasy MinElute spin column, washed, and eluted in RNase-free water. RNA quantity and purity were assessed using a NanoDrop spectrophotometer.

### RNA isolation, cDNA synthesis and RT–qPCR

Total RNA from cultured cells was extracted using TRIzol reagent (Thermo Fisher Scientific) according to the manufacturer’s protocol. Briefly, 5 × 10⁶ cells were homogenized in 500 µl TRIzol, followed by the addition of 100 µl chloroform. Samples were centrifuged at 12,000 × g for 15 min at 4 °C, and the aqueous phase containing RNA was transferred to a fresh tube. RNA was precipitated with an equal volume of 100% isopropanol, washed once with 75% ethanol, air-dried, and dissolved in DEPC-treated H_2_O.

For cDNA synthesis, 1 µg of total RNA was reverse-transcribed using the ABScript II cDNA First-Strand Synthesis Kit (ABclonal) according to the manufacturer’s instructions. The reaction mix contained Random Primer Mix, 10 mM dNTPs, and ABScript II reverse transcriptase. Reactions were incubated at 25 °C for 5 min, followed by 42 °C for 1 h, and terminated at 80 °C for 5 min. Real-time quantitative PCR (RT–qPCR) was performed using Genious 2× SYBR Green Fast qPCR Mix (ABclonal) on a CFX96 Touch Real-Time PCR Detection System (Bio-Rad). Each reaction contained 10 ng of cDNA and gene-specific primers targeting mFwe1, mFwe2, mFwe3, mFwe4, Caspase3, Bax, Bcl2, Gfap, Lcn2, C3, Serping1, S100a10, Ptx3, Megf10, Mertk, Lrp1, and Abca1. GAPDH served as the internal control. Gene expression was normalized to GAPDH using the ΔCt method, and relative expression was calculated by the ΔΔCt method.

### Oligonucleotides

Quantitative PCR (qPCR) was performed using gene-specific primers designed for mouse Flower isoforms (mFwe1–4), apoptosis-related genes, astrocytic markers, inflammatory genes, and lipid metabolism-related genes. Primer sequences were as follows: For mFwe1, forward 5′-ACTGACTCATCCAATAACAGG-3′ and reverse 5′-ACCATGCCTACACATACATTAG-3′; for mFwe2, forward 5′-GCCATTTCTTATGCTCGGAT-3′ and reverse 5′-GCCATTTAACAGAGCTGACA-3′; for mFwe3, forward 5′-GTCTTCTACTGCGGGCAT-3′ and reverse 5′-AGAGGGAAATGGTGTTTCTG-3′; for mFwe4, forward 5′-TTGCTAAATCCTGGGTGTCC-3′ and reverse 5′-TGTTCAGAGGGTGGATAGTG-3′. For apoptosis-related genes, primers were: Bax, forward 5′-CTACAGGGTTTCATCCAG-3′ and reverse 5′-TATTGCTGTCCAGTTCATCT-3′; Bcl2, forward 5′-TACCTGAACCGGCATCT-3′ and reverse 5′-CAGCCAGGAGAAATCAAACA-3′; and Casp3, forward 5′-GGCTTGCCAGAAGATAC-3′ and reverse 5′-CTCCAGGAATAGTAACCAG-3′. For astrocytic and inflammatory markers, primers were: Gfap, forward 5′-AAATTGCTGGAGGGCGAAGAA-3′ and reverse 5′-TGTGAGCCTTTTGAGAGGTCTT-3′; Lcn2, forward 5′-AATGTCACCTCCATCCTGGTC-3′ and reverse 5′-GCCACTTGCACATTGTAGCTC-3′; C3, forward 5′-TCCCAGCTACTAGTGCTACTG-3′ and reverse 5′-CAGTACGATGGTCTCTTCGC-3′; Serping1, forward 5′-AGTGCCCATGATGAGTAGCG-3′ and reverse 5′-CACGGGTACCACGATCACAA-3′; S100a10, forward 5′-GACAAAGGAGGACCTGAGAGTG-3′ and reverse 5′-CTCTGGAAGCCCACTTTGCCAT-3′; and Ptx3, forward 5′-CAATGGACTTCATCCCACCGA-3′ and reverse 5′-ACAGCTTGTCCCACTCCGA-3′. For phagocytosis- and lipid metabolism-related genes, primers were: Megf10, forward 5′-CAGCGCCTATGATGGGGAAT-3′ and reverse 5′-CACTGCCCCGAGATATGGTC-3′; Mertk, forward 5′-CGCTCAGACAATGGGTCGT-3′ and reverse 5′-AAGGCTGTGTTTCTGGTGACA-3′; Lrp1, forward 5′-GTATTGAGCGGTCTCGCCTT-3′ and reverse 5′-CATCCGAGCATCACACCAGT-3′; and Abca1, forward 5′-TAGCAGCACCGTGTCTTGTC-3′ and reverse 5′-GCGTGTCACTTTCATGGTCG-3′. Gapdh was used as the internal reference gene, with forward primer 5′-GTATGTCGTGGAGTCTACTG-3′ and reverse primer 5′-TATTTCTCGTGGTTCACACC-3′.

### Flower topology experiments

To investigate the membrane topology of the Flower protein, mFwe1 and mFwe2 fusion constructs were generated by incorporating the ratiometric pH-sensitive fluorophore pHluorin2. The pHluorin2 tag was fused to either the N- or C-terminus of the Flower protein. Additionally, syb2-pHluorin2 and cytosolic pHluorin2 constructs were used as internal controls. WT astrocytes were transfected individually with each pHluorin2 construct. Following transfection, cells were plated onto glass coverslips and allowed to adhere for 24 h. Fluorescence measurements were acquired using a confocal microscope (LSM 780, Zeiss) equipped with a 40× objective. Images were recorded in 0.5 Hz. Fluorescence was monitored before and after perfusion with 200 µM carbonyl cyanide m-chlorophenyl hydrazone (CCCP, Sigma), using excitation wavelengths of 405 nm and 488 nm. CCCP induces cytosolic acidification, which alters the fluorescent properties of pHluorin2. In acidic conditions, pHluorin2 exhibits a decrease in excitation at 395 nm and an increase at 475 nm.

### Software and Statistical analysis

Image analysis was performed using Fiji (RRID: SCR_002285). Statistical analyses were conducted using SigmaPlot v14.5 and Igor Pro 6.37. Flow cytometry data were analyzed with FlowJo. qPCR data were processed using Bio-Rad CFX Manager software. Figures were prepared using BioRender and CorelDRAW Graphics Suite. The specific statistical tests used are indicated in each figure legend. Student’s t-test, one-way ANOVA or two-way ANOVA were applied, as appropriate, followed by Tukey’s, Dunn’s or Holm–Sidak post hoc tests for multiple comparisons to assess statistically significant differences between groups. All values are presented as mean ± SEM from independent experiments.

## Data Availability

All data supporting the findings of this study are available within the paper and its Supplementary Information file. Raw data and statistical analysis data are provided as a source data file.

## Expanded View Figure Legend

**Figure EV1.**
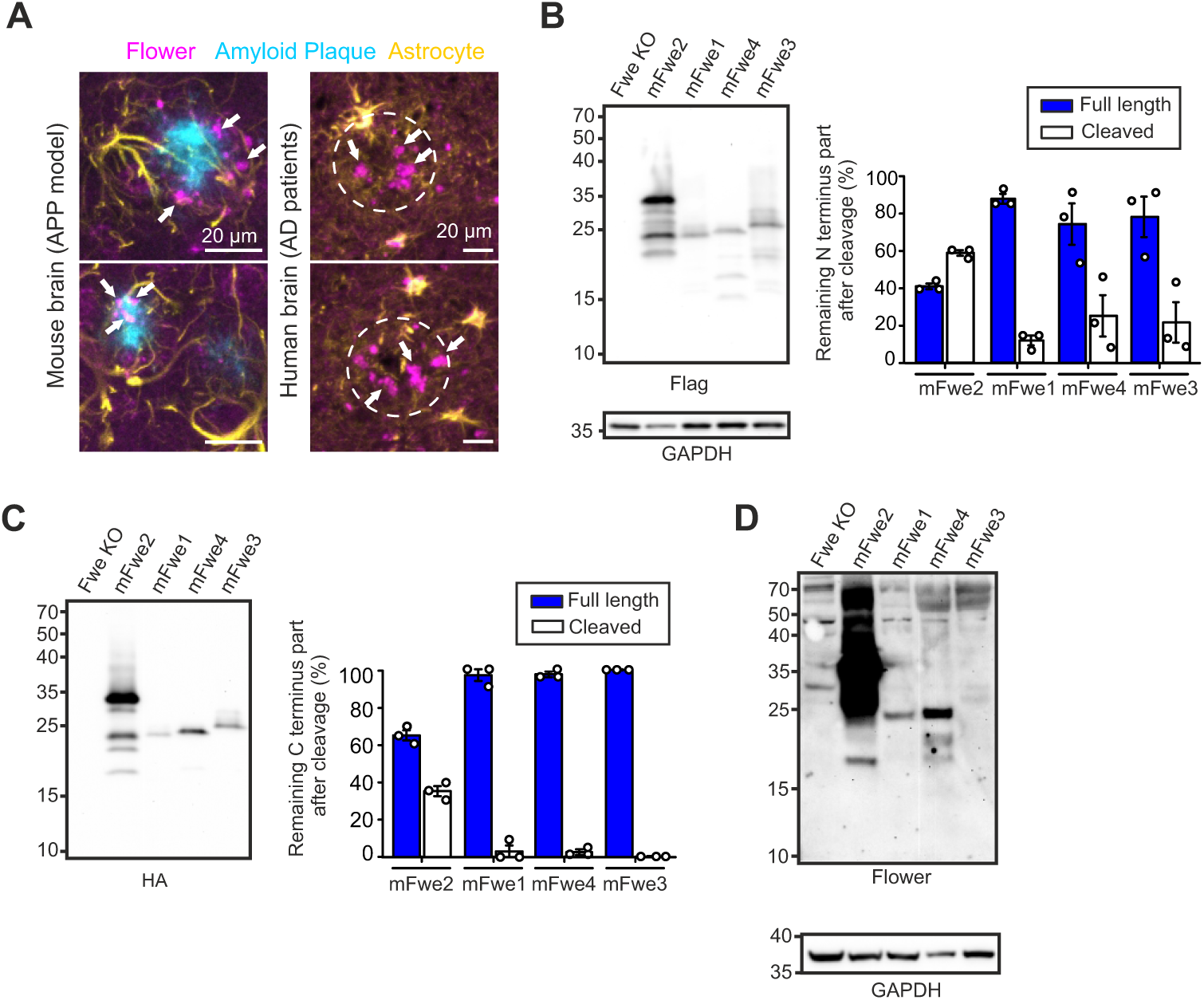
Cleaved Flower fragments in astrocytes, related to Figure 3. **(A)** Confocal images of cortical brain slices from APP-transgenic mice and postmortem human brain tissue from AD patients. Flower isoforms were detected using a C-terminal-specific Flower antibody (#C-1557). Arrows indicate Flower signal localized in the extracellular matrix. A white dashed circle marks a plaque in the human tissue. Scale bar, 20 µm. **(B-C)** Western blot analysis showing proteolytic cleavage of overexpressed Flower isoforms. Flower KO astrocytes were transfected with 3×Flag-mFwe-3×HA constructs, containing a Flag tag at the N-terminus and an HA tag at the C-terminus. Lysates from untransfected Flower KO astrocytes served as negative controls. GAPDH was used as a loading control (N = 3 independent experiments). **(B)** Left: Blot probed with anti-Flag antibody to detect N-terminal fragments from all expressed isoforms. Right: Quantification of full-length and cleaved fragments. **(C)** Left: Blot probed with anti-HA antibody to detect C-terminal fragments. Right: Quantification of full-length and cleaved fragments. **(D)** Longer exposure of a separate blot from the same experiment, reprobed with anti-Pan-Flower antibody to confirm band identity.

**Figure EV2.**
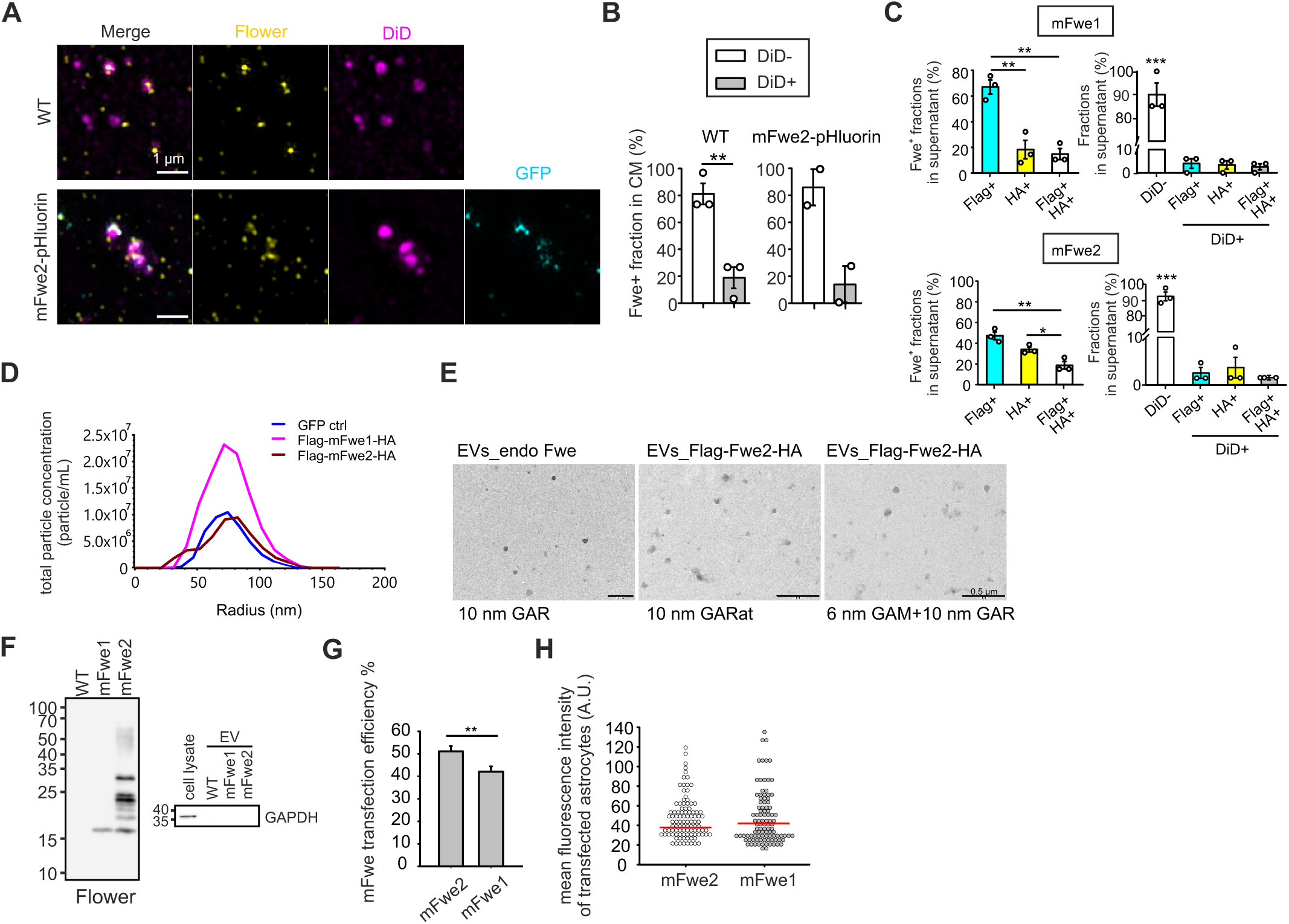
Isoform-specific association of Flower with extracellular vesicles in astrocytes, related to Figure 3. **(A)** SIM images of supernatant fractions collected from WT or mFwe2-pHluorin2-expressing astrocyte cultures. Pelleted fractions were deposited onto poly-ornithine-coated glass coverslips, fixed, and stained with a polyclonal Pan-Flower antibody recognizing both N- and C-termini, together with DiD to label lipid membranes. In mFwe2-overexpressing cultures, anti-GFP antibody was used to detect overexpressed Flower. Scale bar, 1 µm. **(B)** Quantification of co-localization between Flower-positive particles and the lipid dye DiD from (**A**). Statistical significance was determined using Student’s t-test: **p < 0.01. **(C)** Analysis of secreted Flower from overexpressing astrocytes shown in Fig. 3B. Supernatants from Flower KO astrocyte cultures transduced with 3×Flag-Flower-3×HA constructs were analysed by immunostaining with anti-Flag (N-terminal) and anti-HA (C-terminal) antibodies. Co-staining with a lipid dye assessed potential association with EVs. SIM was performed on supernatants from mFwe1- and mFwe2-expressing cells (N = 3; mFwe1: 5,330 particles; mFwe2: 5,348 particles). Data are presented as mean ± SEM. Statistical significance was determined by one-way ANOVA followed by Tukey’s post hoc test (*p < 0.05, **p < 0.01). **(D)** Particle analysis of EVs isolated from astrocyte cultures expressing GFP vector control, Flag-mFwe1-HA, or Flag-mFwe2-HA. Particle number and radius are shown. **(E)** Background control for immunogold labeling experiments shown in Fig. 3C and D. Immuno-labeling was done under the same conditions as described in Fig. 3C and D, without the application of a primary antibody to confirm the gold bead background. From left to right: electron micrographs of EVs immune-labeled with goat anti-rabbit (10 nm, GAR), goat anti-rat (10 nm, GARat) and goat anti-mouse (6 nm, GAM) together with GAR (10 nm) are shown. Scale bar, 0.5 µm. **(F)** Western blot analysis of secreted EV lysates from WT astrocyte cultures transfected with mFwe1 or mFwe2. Ten micrograms of EV protein were loaded per lane. The blot shown in Fig. 3G was stripped and sequentially reprobed with anti-Pan-Flower antibody (left). Anti-GAPDH antibody was used to assess EV purity (total astrocyte lysate included as a positive control) (right). **(G)** Primary mouse astrocytes were transfected with either the “lose” isoform (mFwe1-mCherry or mFwe1-mTFP) or the “win” isoform (mRFP-mFwe2 or mFwe2-mTFP) to assess transfection efficiency. The number of fluorescent mFwe1⁺ and mFwe2⁺ cells was quantified from confocal images of separately cultured populations. N = 2 independent preparations; n = 84 cells for mFwe1, n = 101 cells for mFwe2. Statistical analysis was performed using Welch’s t-test; two-tailed P < 0.001 (**). **(H)** Mean fluorescence intensity per cell from the same dataset as in (**G**), used to assess expression levels of mFwe1 and mFwe2. Red lines indicate median values. N = 2 independent preparations; n = 100 cells per group. Statistical comparison using the Mann–Whitney U test showed no significant difference.

**Figure EV3.**
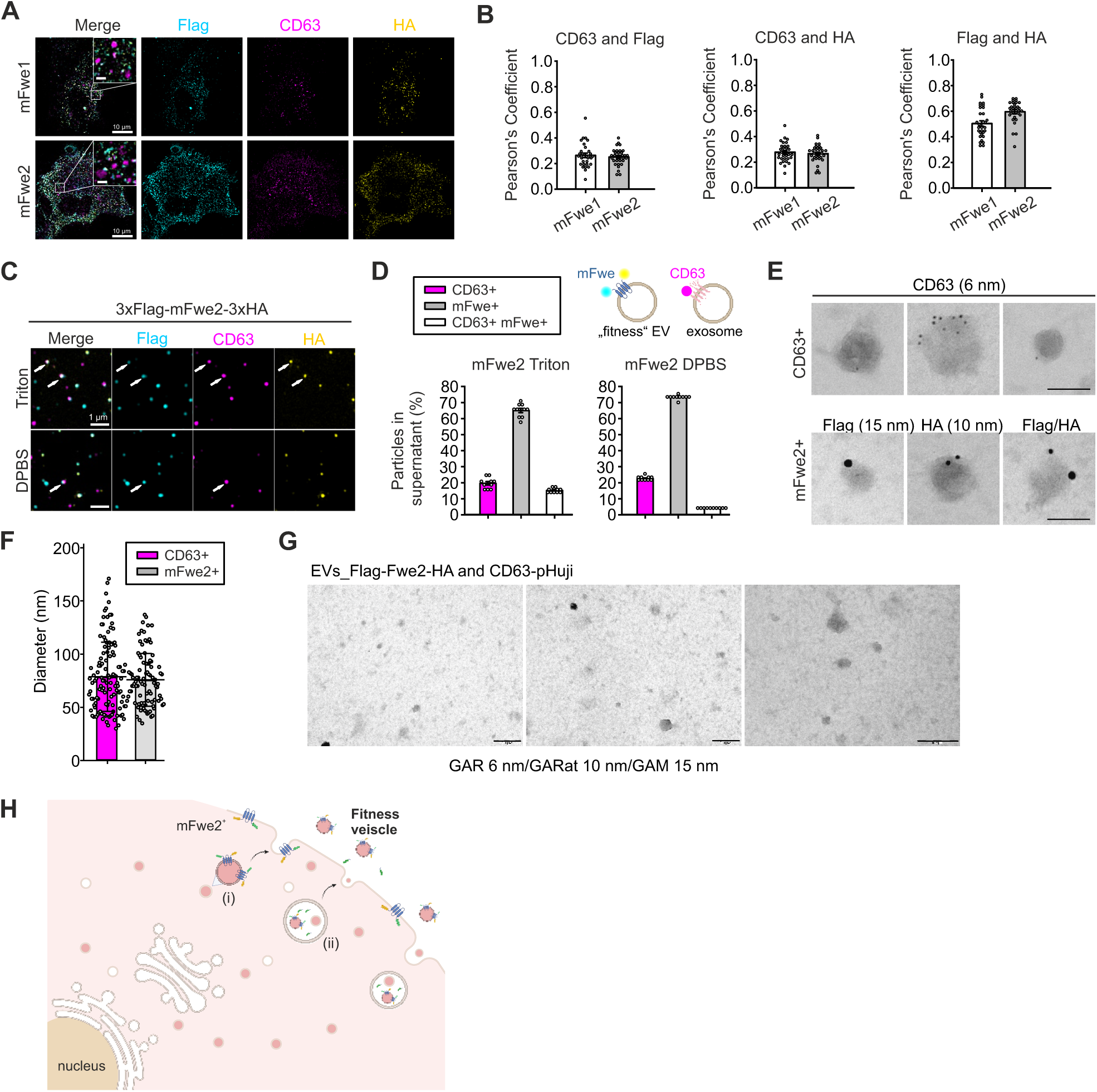
Flower-positive EVs represent a secretory population distinct from CD63⁺ exosomes, related to Figure 3. **(A)** SIM images of WT astrocytes co-transfected with CD63-pHuji together with mFwe1 or mFwe2 (3×Flag-Flower-3×HA). Cells were fixed 24 h after transfection and stained with anti-RFP, anti-Flag and anti-HA antibodies. Scale bar, 10 µm; inset: 1 µm. **(B)** Quantification of colocalization between CD63-phuji and Flag- or HA-tagged Flower isoforms from (**A**), expressed as Pearson’s correlation coefficient. N = 4 pups for mFwe2 and N = 2 pups for mFwe1; n = 30 cells analyzed per group. **(C)** SIM images of WT astrocytes co-transfected with CD63-pHuji and mFwe2 (3×Flag–Fwe2–3×HA). EVs were harvested 3 days after transfection, fixed, and stained with anti-RFP, anti-Flag, and anti-HA antibodies under permeabilized or non-permeabilized conditions, as indicated. White arrows indicate CD63⁺/mFwe⁺ EVs. Scale bar: 1 μm. **(D)** Quantification of the percentage of CD63-positive puncta and mFwe-positive puncta under permeabilized and non-permeabilized conditions, analyzed from (**C**). HA single-positive, Flag single-positive, and double-positive puncta were defined as mFwe events. N = 4 pups for mFwe2 and N = 2 pups for mFwe1; n = 10 images analyzed per group. Upper Right: Schematic illustrating the transmembrane topologies of mFwe-bearing EVs and CD63-pHuji-labeled exosomes. **(E)** Representative electron micrographs of EVs used from (**C**). Co-immunogold labeling was performed using primary antibodies against pHuji (anti-mRFP), Flag, and HA, detected with goat anti-rabbit (6 nm), goat anti-mouse (15 nm), and goat anti-rat (10 nm) gold-conjugated secondary antibodies, respectively. Scale bar, 100 nm. **(F)** Size distribution analysis of immunogold-labeled EVs shown in (**E**). Data are presented as mean ± SD. Anti-CD63⁺ EVs, n = 118; anti-HA-, anti-Flag-, or double-positive EVs (mFwe⁺), n = 94; N = 4 pups. **(G)** Background control for immunogold labeling experiments shown in (**E**). Immuno-labeling was done under the same conditions as described in (**E**), without the application of primary antibodies to confirm the gold particle background. Scale bar, 0.2 µm. **(H)** Model of heterogeneous Flower-containing vesicle populations in astrocytes. Schematic illustrating the two primary fates of Flower protein during astrocyte secretion. (i) Surface-retained Flower: Following vesicle fusion, a population of Flower protein remains integrated into the plasma membrane. (ii) Secreted Flower-EVs: Flower is released within a distinct class of extracellular vesicles. In both populations, the N- and C-termini are oriented toward the extracellular space (“extracellular-out”), rendering them accessible for intercellular interactions.

**Figure EV4.**
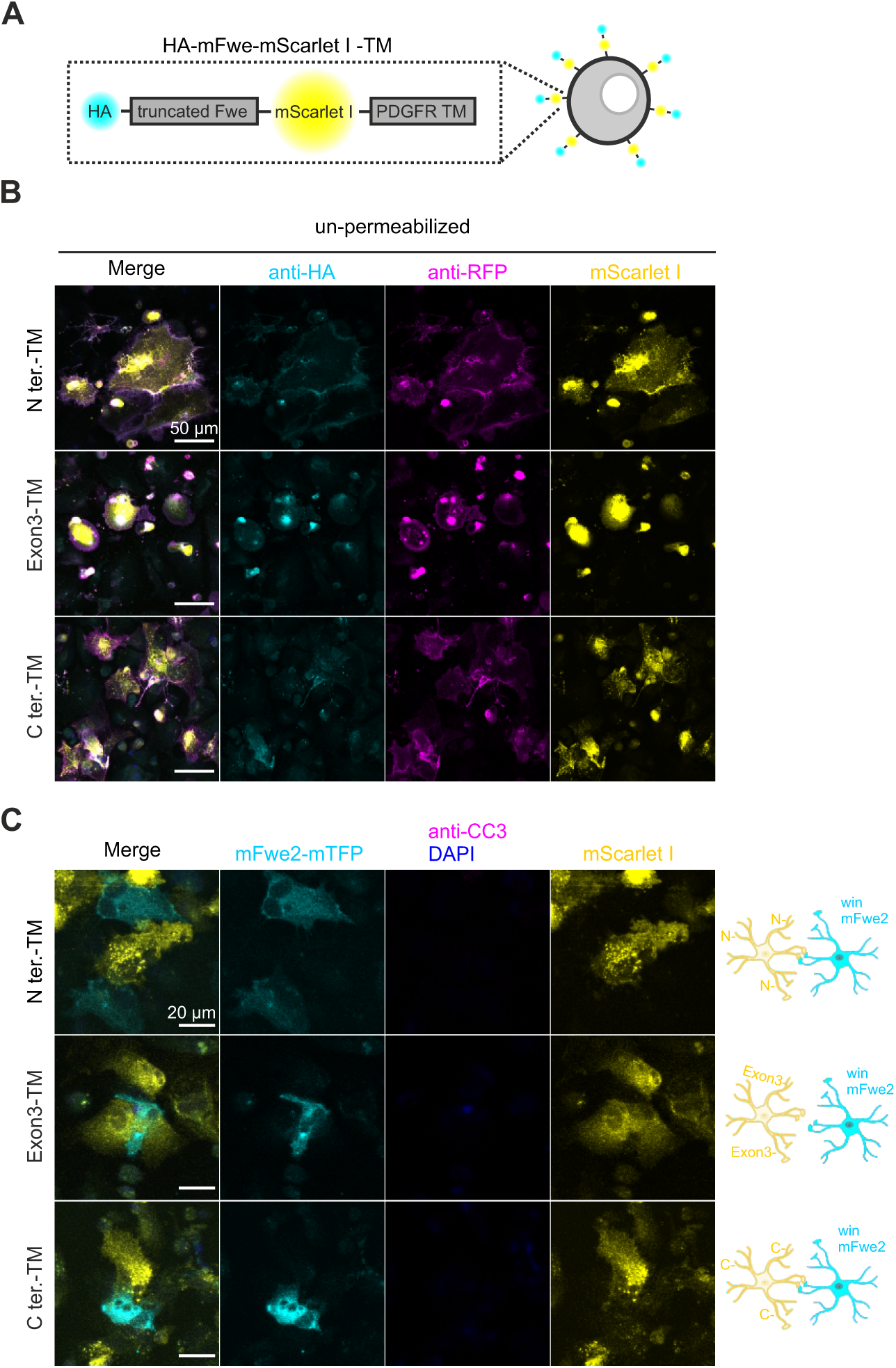
The N-terminal Flower domain does not induce cell death in fit cell populations, related to Figure 4. **(A)** Schematic of constructs in which truncated Flower domains (N-terminus, Exon 3, or C-terminus) were fused to a transmembrane (TM) domain for plasma membrane (PM) localization with extracellular orientation. An HA tag was placed at the N-terminus and mScarlet I at the C-terminus to assess membrane orientation. **(B)** Confocal images of Flower KO astrocytes expressing truncated Flower domains targeted to the PM. To verify surface expression, fixed, non-permeabilized cells were stained with anti-RFP-Alexa647 (magenta) and anti-HA (cyan). Scale bar, 50 µm. **(C)** Confocal images of Flower KO astrocytes expressing specific truncated Flower domains (yellow) co-cultured with fit mFwe2-mTFP-expressing “win” cells (cyan). Cultures were stained with DAPI before fixation and with anti-CC3 (magenta) after fixation to detect apoptotic cells. Maximum intensity projection images are shown. Scale bar, 20 µm.

**Figure EV5.**
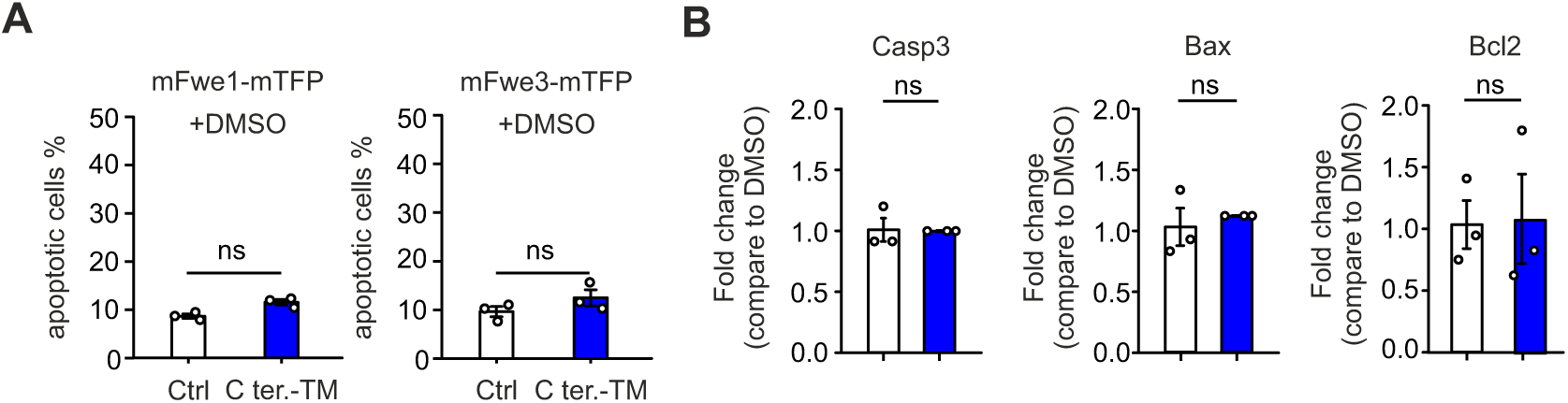
Overexpression of the C-terminus Flower domain does not alter cell survival, related to Figure 4. **(A)** Flower KO astrocytes were transfected with mFwe1 or mFwe3 to generate unfit cell populations. Co-transfection with the C-terminal Flower domain (C-term-TM) plasmid did not affect cell survival upon treatment with the control vehicle, DMSO. **(B)** Quantitative RT-PCR analysis of apoptosis-related genes in transfected and treated cells. N = 3 independent experiments. Data are presented as mean ± SEM. Statistical significance was determined using Student’s t-test: ns refers to not significant.

**Figure EV6.**
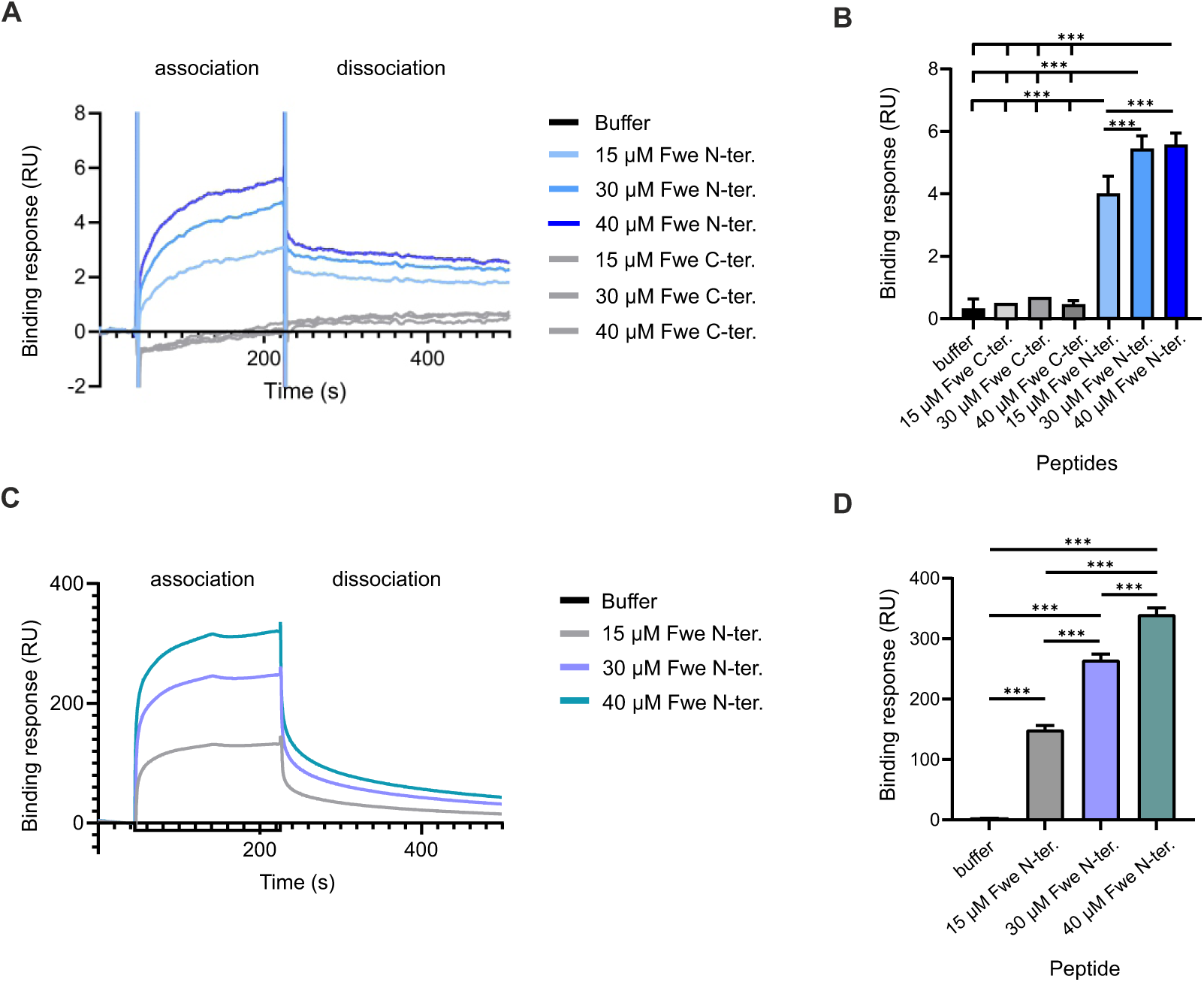
SPR analysis showing interaction between the N- and C-terminal and N- and N-terminal domains of Flower, related to Figure 5. **(A-B)** SPR binding analysis showing interaction of Flower N-terminal and C-terminal peptides with immobilized Flower C-terminal peptide (engineered with an N-terminal cysteine for immobilization). **(C-D)** SPR binding analysis showing interaction of Flower N-terminal and N-terminal peptide with immobilized Flower N-terminal peptide. Representative sensorgrams **(A, C)** and the corresponding quantification **(B, D)** demonstrate binding responses at increasing concentrations (15, 30, and 40 μM). Bar graph shows mean ± s.d. from three independent experiments. One-way ANOVA: **p < 0.01, ***p < 0.001.

**Figure EV7.**
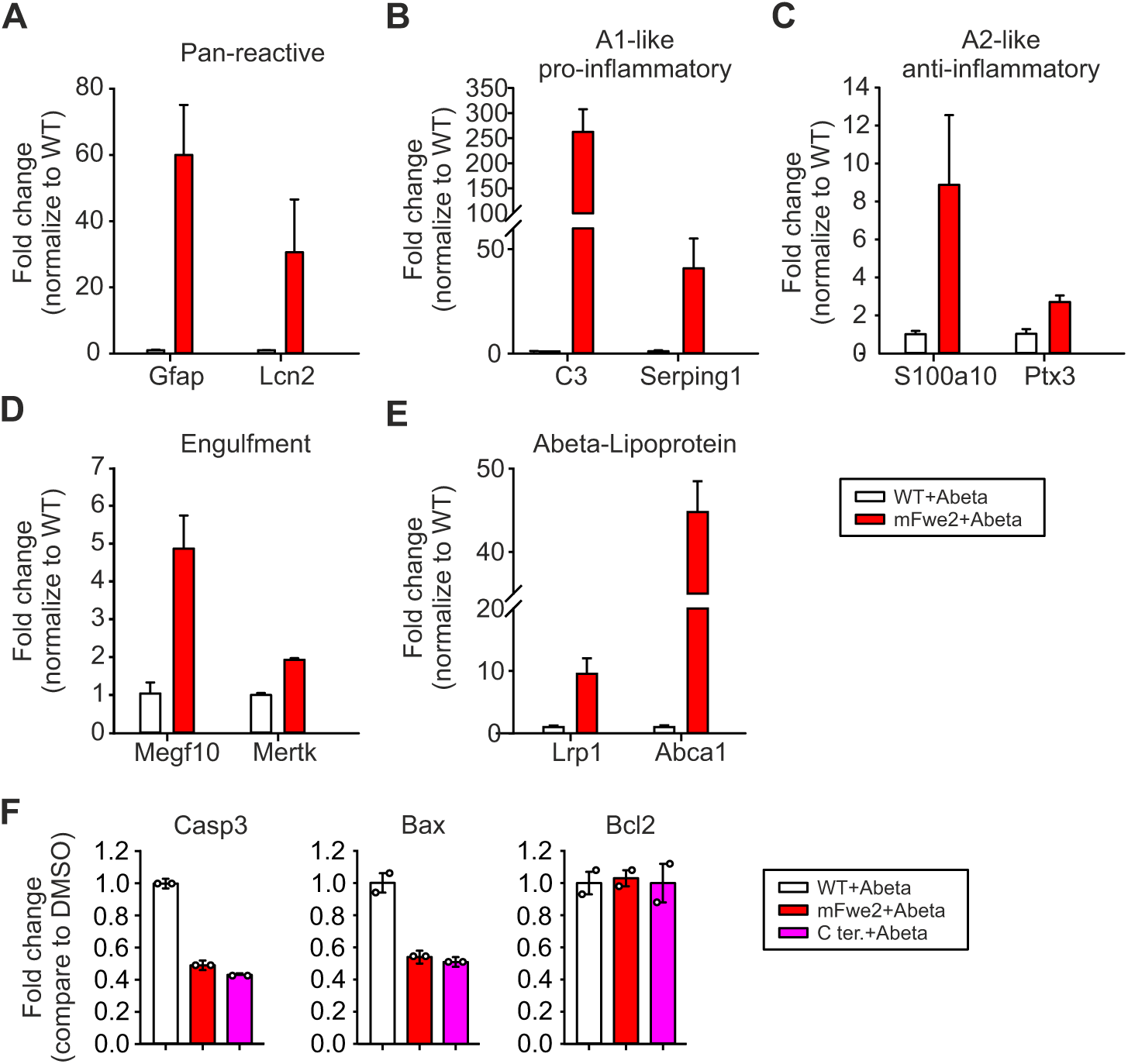
Overexpression of Flower upregulates genes associated with astrocyte reactivation and lipid transport in an Aβ protofibril environment, related to Figure 7. WT astrocytes and astrocytes lentivirally infected with mFwe2 construct were co-cultured with Aβ protofibrils for six days. **(A-E)** RNA was isolated from each group and analyzed by quantitative RT–PCR for the indicated genes. The indicated genes are shown. **(F)** Quantitative RT–PCR analysis of apoptosis-related genes (*Caspase-3, Bax, and Bcl2*) (N = 2 independent experiments). Data are presented as mean ± SEM.

## ACKNOWLEDGEMENTS

This work was supported by grants from the Deutsche Forschungsgemeinschaft (DFG), collaborative research center SFB894 (ID number 157660137), subproject A10 (to H.-F.C. and E.K.), and A14 (to V.F.) and FL153/10-2 (to V.F.), and the University of Saarland (HOMFORexzellent 2020, NanoBioMed Young Investigator Grant 2020, and HOMFOR Anschubfinanzierung 2025 to H.-F.C.). The surface plasmon resonance instrument was acquired for the Center for Molecular Signaling (PZMS) according to Art. §91b GG (SL1369001; INST 256/548-1). Part of the laser capture microdissection equipment was funded by the German Research Foundation (DFG; ID number 447452855; INST 256/541-1). Furthermore, we thank Keerthana Ravichandran, Margarete Klose, Anja Bergsträßer, Nicole Rothgerber, Katrin Sandmeier, Andrea Schottek and Christine Wesely for their excellent technical assistance.

## AUTHOR CONTRIBUTIONS

H.-F.C. conceived and supervised the study. S.-M.T. performed all cellular fitness assays, molecular biology experiments, and tissue staining analyses. C.-H.L. and Y.S. assisted with neuronal culture preparation. L.Y. carried out AD human brain tissue sectioning and staining. J.H. performed laser microdissection of brain tissue. M.J. synthesized the peptides. C.S. performed EM analysis A.C. and S.R. carried out the SPR analysis. A.A. and D.Y. performed particle size analysis. W.J.S.-S. provided postmortem AD brain tissue. C:-A.Y. conducted Flower gene expression analysis in AD cohorts. V.F. generated the Flower antibodies and Flower-gene-deficient mice. H.-F.C. and E.K. secured funding for the project. L.Y. and C.-K.Y. and C. S. contributed essential intellectual input to the study. C.-K.Y., C.S. and V.F. critically reviewed the manuscript. H.-F.C. wrote the manuscript with input from all authors. All authors reviewed and approved the final version of the manuscript.

